# Computational Transcriptomic and Comparative Genomic Analysis of dead box RNA Helicase gene AT2G45810 Expressed in Plants Arabidopsis thaliana

**DOI:** 10.1101/2024.01.12.575476

**Authors:** Mohammed Emon, Akram Hosen, Sujay Kumar Bhajan, Zilhas Ahmed Jewel, Md. Sarafat Ali

## Abstract

*Arabidopsis thaliana* is a short life cycle, small genome, and Brassicaceae family winter annual small flowering plants. It is popularly used as a model organism in genetics and plant biology research, and it is essential to understanding the molecular biology of many plant features, including light sensing and flower formation. It also plays a key role in the science of agronomy, and plant transcriptomics as well as genomics. When it comes to the development of multicellular creatures, transcriptional programs are crucial. The constantly active growth of different organ systems is supported by transcriptional programs. Arabidopsis embryos possess remarkable transcriptomes compared to other plant tissues comprising somatic embryo differentiation circumstances operating during plant embryogenesis. Here we show that the transcriptomic analysis of the genome dead box RNA helicase gene AT2G45810 of *Arabidopsis thaliana* Araport11 species which revealed the specific gene expression patterns of Arabidopsis tissue-specific information of developmental map, embryo, single cell, DNA damage, cellular interactions, pathway analysis, etc., through *In Silico* or computational approaches. In this particular study, we used the TAIR, Phytozome, and plant comparative genomics portal for retrieving and identification of specific genes of interest. Next, we used web-based Bar utoronto tools to visualize other data, including functional genomics. Their protein and gene expression tools facilitate the exploration of promoters, the identification of protein-protein interactions, the viewing of expression patterns as electronic fluorescent pictographs or heatmaps, and more.

## Introduction

*Arabidopsis thaliana,* a winter annual exhibiting a comparatively short lifecycle, is a widely utilized model organism in the field of plant biology and genetics. *A. thaliana*, being a complex multicellular eukaryote, possesses a genome size of approximately 135 mega base pairs(1,2). *Arabidopsis thaliana,* commonly known as *A. thaliana,* has emerged as a prominent model organism in the field of plant sciences. Its utilization encompasses various areas such as genetics, evolution, population genetics, and plant development. *A. thaliana*, despite its limited agricultural relevance, possesses various characteristics that render it a valuable model organism for comprehending the genetic, cellular, and molecular intricacies of flowering plants(3–5).

*Arabidopsis thaliana* is a comparatively minute plant in the mustard family that has established the reference plant for research in plant biology favorable for genetic and molecular research(6,7). More vigorously, it can also provide a compatible model for plant biology along with recognizing rudimentary queries of biological conformation with functions common to all eukaryotes(6). Rapid, inexpensive, and expedient research can be adopted using *Arabidopsis thaliana*(8). The plants grow from a seed to a plant producing mature seeds in as bit as 6 weeks, relying on growth conditions(6,8). In the laboratory, it can grow easily under the condition of dim fluorescent lighting that is facile to achieve indoors(6,8,9). Not only it has small seeds and seedlings which can also grow on a single Petri dish. Some exceeding features like co-culture are not necessary, supporting aseptic growth conditions, and huge control in variability(6,8,9). *Arabidopsis thaliana* can accumulate both positively and negatively charged nano plastics which are promoted by the growth medium and root exudates. Thus, positively charged nano plastics accumulated at relatively low levels in the root tips the positively charged nano plastic deposited negatively charged nano plastics were found repeatedly in the apoplast and xylem(10).

*Arabidopsis thaliana* has many amenities for genome study, such as a short generation period, minute size, a large number of offspring, and a relatively small nuclear genome(6,7,11). The growth of the scientific community has influenced the advantages of these plants(7,12). The roughly 146 million base pairs that make up the haploid genome of *Arabidopsis thaliana* are spread across five different chromosomes. Of these, 25 000 projected genes that encode proteins are present, and almost 85% of these genes are part of families with two to several hundred members as well as 38,000 loci(12,13). The sequenced regions screen 115.4 megabases of the 125-megabase genome and extend into centromeric regions even 30 megabases of the annotated genomic sequence have already been entrusted to GenBank(6,12).

By using the study of phenotype and genotype can analyze the functions because of containing lots of collection of gene disruptions and the presence of the entire genomic sequence(7,12). By genomic analysis, can be possible to provide several methods and resource materials including easy procedures for chemical and insertional mutagenesis, efficient mechanisms for performing crosses and introducing DNA through the transformation of plants, grant addition of mutants with distinct phenotypes, and a variety of chromosome maps(6).

In Arabidopsis having forward-genetic screening developed mutagenized seeds as well as ethyl methane sulfonate, irradiation, bombardment, and sodium azide, have coincided with the nonplant model system on Arabidopsis. Plants are exposed to immunity by many pathogenic organisms, able to recognize and respond to the plethora of challenges in the case of recognition, and act a variety of cell-surface-localized pattern recognition receptors (PRRs). These receptor patterns recognize pathogen-associated molecular patterns (PAMPs), and danger-associated molecular patterns (DAMPs) as well as induce pattern-triggered immunity (PTI), and some of the patterns were used in two-week-old *Arabidopsis thaliana* for four replicate experiments(14–16). It is possible to target mutagenesis and update using CRISPR/Cas9 gene editing(8,17). Transcriptional programs underlie the development of various organ systems therefore Arabidopsis embryos possess remarkable transcriptomes compared to other plant tissues comprising somatic embryo differentiation and lead embryogenesis(18).

The European Space Agency is conducting ongoing research on A. thaliana aboard the International Space Station. The study of plant growth and reproduction in microgravity, from seed to seed, is the main objective(19,20).

Here we show the transcriptomic and comparative genomics analysis of the genome dead box RNA helicase of *Arabidopsis thaliana* Airport11 (AGI ID: AT2G45810) species revealed the specific gene expression patterns of Arabidopsis tissue-specific information of developmental map, embryo, single cell, DNA damage, cellular interactions, subcellular localization, pathway visualization and analysis, promoter analysis and so on.

## 2. Methods and Materials

### 2.1 Data retrieval and sequences analysis

Gene sequence information was accessed from Araport(21,22) at https://www.araport.org/. We retrieved the *Arabidopsis thaliana* gene ID by using Phytozome(23) (https://phytozome-next.jgi.doe.gov/) as selecting genome *Arabidopsis thaliana* Araport11-thale cress and gene-specific dead box RNA helicase(24).

**Fig 1.**
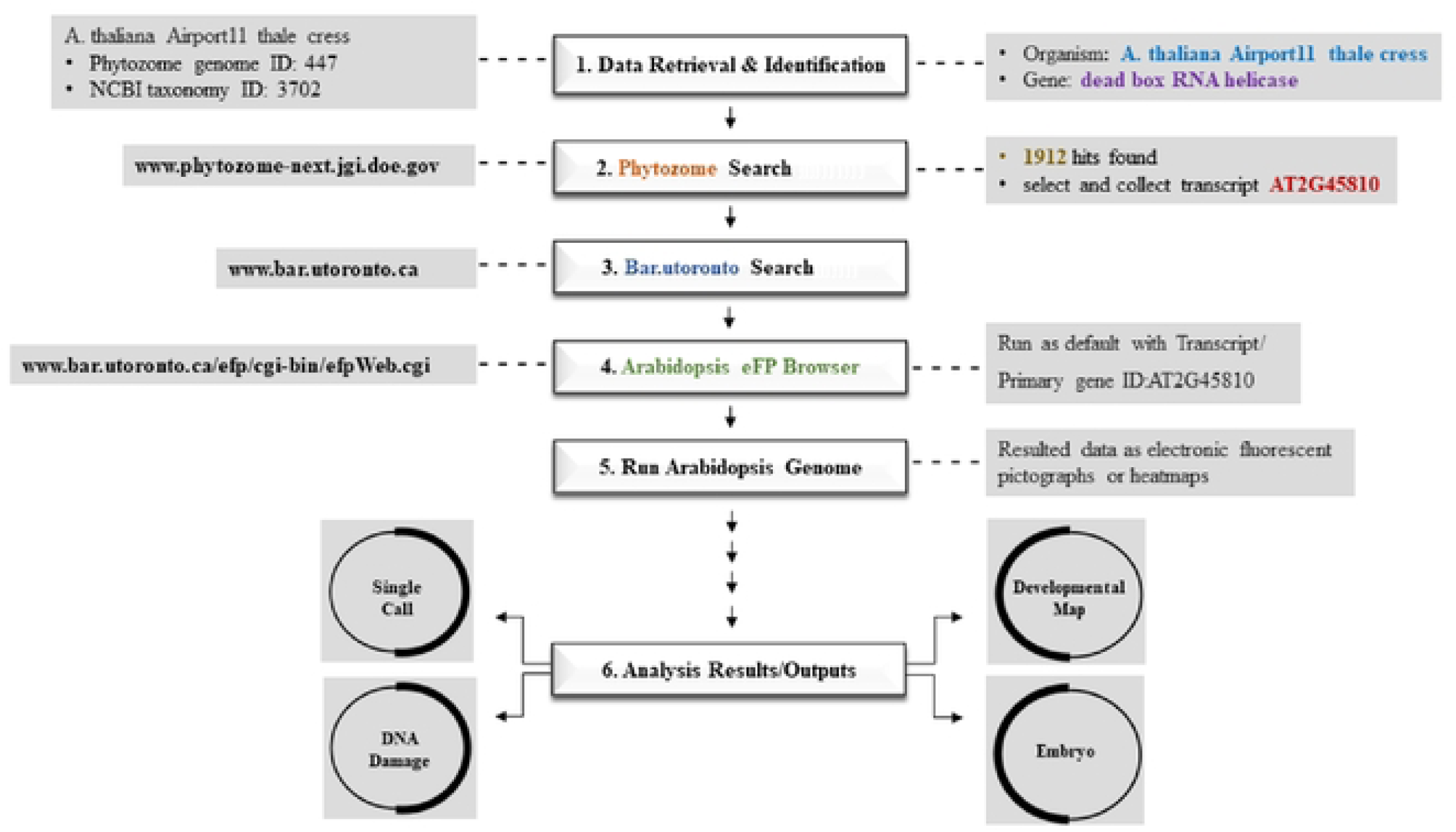
Flowchart: A general sketch of the study integrated with the use of several computational tools and their approaches to computational transcriptomic and comparative genomic analysis of gene AT2G45810 expressed in the Arabidopsis thaliana.

### 2.2 Phylogenetics analysis

A custom phylogenetic tree was constructed using AT2G45810 and its paralogs in Arabidopsis to corroborate the curated trees using PLAZA 5.0(25) with the 9 Blast hits identified for *A. thaliana* in Data Settings → Species Selection, with other parameters as default (https://bioinformatics.psb.ugent.be/plaza/versions/plaza_v5_dicots/analysis/custom_tree_creation/gene_id/AT2G45810.

### 2.3 Subcellular localization and significance analysis

Subcellular localization of this set of genes was identified using SUBA5 at https://suba.live/suba-app/factsheet.html?id=AT2G45810.1,(26,27) And the significance of these localizations was evaluated with the Cell eFP Browser(28) at http://bar.utoronto.ca/cell_efp/cgibin/cell_efp.cgi.

### 2.4 Expression analysis

The tissues in which AT2G45810 and its 5 in-paralogs were highest expressed were identified across the different compendia (i.e. Data Sources) available in the Arabidopsis eFP Browser(28) at http://bar.utoronto.ca/efp/cgi-bin/efpWeb.cgi. TraVA(29) at http://travadb.org/ using the TMM normalization type and JBrowse from Araport(22) at https://apps.araport.org/jbrowse/?data=arabidopsis viewing available read alignment tracks were used to corroborate the results from the eFP Browser.

### 2.5 Co-expression analysis

In the computational tool BAR utoronto’s Expression Angler which help to determine which genes display co-expression, we search to identify genes based on the recurring nature of gene AT2G45810 expression. Searching as step 1, we have entered AGI ID or gene aliase AT2G45810 in the input box then we selected a view on Development map and Tissue Specific from the list Co-expression gene sets were generated using Expression Angler(30) at https://bar.utoronto.ca/ExpressionAngler/ by selecting the top 50 co-expressed genes of AT2G45810 from the views (compendiums) in which it was most expressed and those in which its homologs were most expressed, which were the Tissue Specific and Developmental Map views, respectively. Furthermore, based on the expression levels of the genes, a “Custom Expression Pattern” was established to find genes in the Tissue-Specific view with expression patterns comparable to AT2G45810. Using the “coex in specific conditions (Ath)” option and sorting on the “MR (tissue)” column, ATTED-II(31) at https://atted.jp/ was used to identify the genes that have the largest mutual rank (MR) with AT2G45810. Employing the Aranet V2(32) program at https://www.inetbio.org/aranet/, functional inference via guilt-by-association was examined. Query option 1: “Find new members of a pathway” was used with default settings. By downloading the output.csv or.txt files from these tools and utilizing the Venn Selector tools, which are accessible at the BAR (http://bar.utoronto.ca/ntools/cgi.bin/ntools_venn_selector.cgi), the overlap of sets was accomplished.

### 2.6 Promoter analysis

To analysis and revealed the promoter analysis of *Arabidopsis thaliana* specific genes we used and examined several manipulated and accepted online tools such as ATTED II, PLACE, Cistome. The 20 “Tissue Specific” co-expressed genes with a Custom Expression Pattern like AT2G45810 from the Expression Angler tool that was in common with the ATTED II (https://atted.jp/coexsearch/) tool’s top 300 “MR (tissue)” “coex in specific conditions (Ath)” gene were investigated for common motifs from the PLACE database using the Cistome tool(30) at https://bar.utoronto.ca/cistome/cgi-bin/BAR_Cistome.cgi. The TAIR Upstream (TSS/TrSS) 1000 bp promoter data set was used, and for step 3 “Paste in your own PSSMs or consensus sequences and/or choose precomputed motifs from various sources for de novo mapping” was used, with “All PLACE elements”. Only Significant Motifs were chosen to be shown, with a Ze cutoff > 2 (instead of the default 3, other defaults were kept as is). The top 50 co-expressed genes from just the Expression Angler “Tissue Specific” output were also investigated.

### 2.7 Functional annotations

Functional annotations was predicted through several online databases i.e. InterPro (https://www.ebi.ac.uk/interpro/), ENZYME (https://enzyme.expasy.org/), QuickGO (https://www.ebi.ac.uk/QuickGO/), KEGG (https://www.genome.jp/kegg/kegg2.html), PANTHER (https://www.pantherdb.org/), PFAM (http://pfam.xfam.org/), SMART (https://smart.embl.de/), PROSITE (https://prosite.expasy.org/), SSF (https://supfam.org/SUPERFAMILY/index.html), CATH (http://www.cathdb.info/search).

### 2.8 Functional classification and pathway visualization

Using the top 50 co-expressed genes that exhibit the same “Tissue Specific” expression pattern as AT2G45810 from Expression Angler tool, enriched GO terms were identified using AgriGO V2(33) (reference genome: Affymetrix ATH1 Genome Array (blast); test: hypergeometric test with Hochberg FDR correction) at http://systemsbiology.cau.edu.cn/agriGOv2/. After experimenting with a few different settings, it was found that the GO Slim analysis produced a set of terms that were too high level to be very informative, so the entire set of GO terms was used (because Provart and Zhu’s Classification Superviewer only uses GO Slim terms, its output was also deemed uninformative). Furthermore, it was determined that a cutoff of mapping to at least three words would yield a more illuminating result than the default of five. By entering the table gene list into server g:Profiler(34) at https://biit.cs.ut.ee/gprofiler/, evidence for the GO annotations enriched in this collection was investigated. To be as near to the AgriGO settings as practicable, with the species selected to be *Arabidopsis thaliana*, the significance threshold set to “Benjamini Hochberg FDR”, the background set to “AFFY_ATH1_121501”, and the statistical domain scope set to “Custom over all known genes” (all other settings were left as their defaults). This gene set’s possible pathways were investigated using Aracyc’s Cellular Overview tool,(35) which can be found at https://pmn.plantcyc.org/overviewsWeb/celOv.shtml.

### 2.9 Protein homologs identification

Protein homologs were predicted through various online databases including NCBI Basic Local Alignment Search Tool (BLAST)(36) (https://blast.ncbi.nlm.nih.gov/Blast.cgi), and JBrowse (https://jbrowse.org/jb2/)(37).

### 2.10 Bio sequence analysis

PlantFAMs Bio sequence of *Arabidopsis thaliana* gene AT2G45810 analysis by using computation tool HMMER web server(38) (https://www.ebi.ac.uk/Tools/hmmer/search/hmmsearch).

### 2.11 Network Tools

Interactions with gene AT2G45810 were investigated with Arabidopsis Interactions Viewer 2 (AIV-2)(39) using default parameters at https://bar.utoronto.ca/interactions2/. Enriched transcription factors for the co-expressor set of the top 50 genes showing similar expression patterns as AT2G45810 were identified with TF2Network(40) at http://bioinformatics.psb.ugent.be/webtools/TF2Network/ using the gene identifiers in Table 2. ePlant(41) and GeneMANIA(42) were also used, with default settings.

## 3. Result and Discussions

### 3.1 Genomics retrieval and identification

We elucidated genomics information from Bar utoronto tool’s Arabidopsis eFP Browser. Several genomics information such as assembly source, version, annotation source, version, number of protein-coding transcripts, number of protein-coding genes, etc., significantly analyzed and enlisted in table 1. total Scaffold length 119,667,750 (bp), number of Scaffolds 7, min. number of Scaffolds containing half of assembly (L50) 3, shortest Scaffold from L50 set (N50) 23,459,830, total contig length (bp) 119,482,012, number of contigs 169, number of protein-coding transcripts 48,456, number of protein-coding genes 27,655, percentage of eukaryote BUSCO genes 98.7, percentage of embryophyte BUSCO genes 99.3 etc.,

**Table 1.** Genome information of Arabidopsis thaliana Araport11.

| Serial | Parameters | Results |
| --- | --- | --- |
| 1 | Assembly Source: | TAIR |
| 2 | Assembly Version: | TAIR9 |
| 3 | Annotation Source: | Araport |
| 4 | Annotation Version: | Araport11 |
| 5 | Total Scaffold Length (bp): | 119,667,750 |
| 6 | Number of Scaffolds: | 7 |
| 7 | Min. Number of Scaffolds containing half of<br>assembly (L50): | 3 |
| 8 | Shortest Scaffold from L50 set (N50): | 23,459,830 |
| 9 | Total Contig Length (bp): | 119,482,012 |
| 10 | Number of Contigs: | 169 |
| 11 | Min. Number of Contigs containing half of<br>assembly (L50): | 5 |
| 12 | Shortest Contig from L50 set (N50): | 10,898,021 |
| 13 | Number of Protein-coding Transcripts: | 48,456 |
| 14 | Number of Protein-coding Genes: | 27,655 |
| 15 | Percentage of Eukaryote BUSCO Genes: | 98.7 |
| 16 | Percentage of Embryophyte BUSCO Genes: | 99.3 |

### 3.2 Phylogenetic analysis

The phylogenetic relationship of the specific gene AT2G45810 *Arabidopsis thaliana* was modified and investigated by the reliable PLAZA server. (https://bioinformatics.psb.ugent.be/plaza/versions/plaza_v5_dicots/) The most significant biological discoveries and a deeper comprehension of the patterns along with processes of evolution can be gained by comparing gene sequences or plant species in a phylogenetic framework(43).

**Fig 2.**
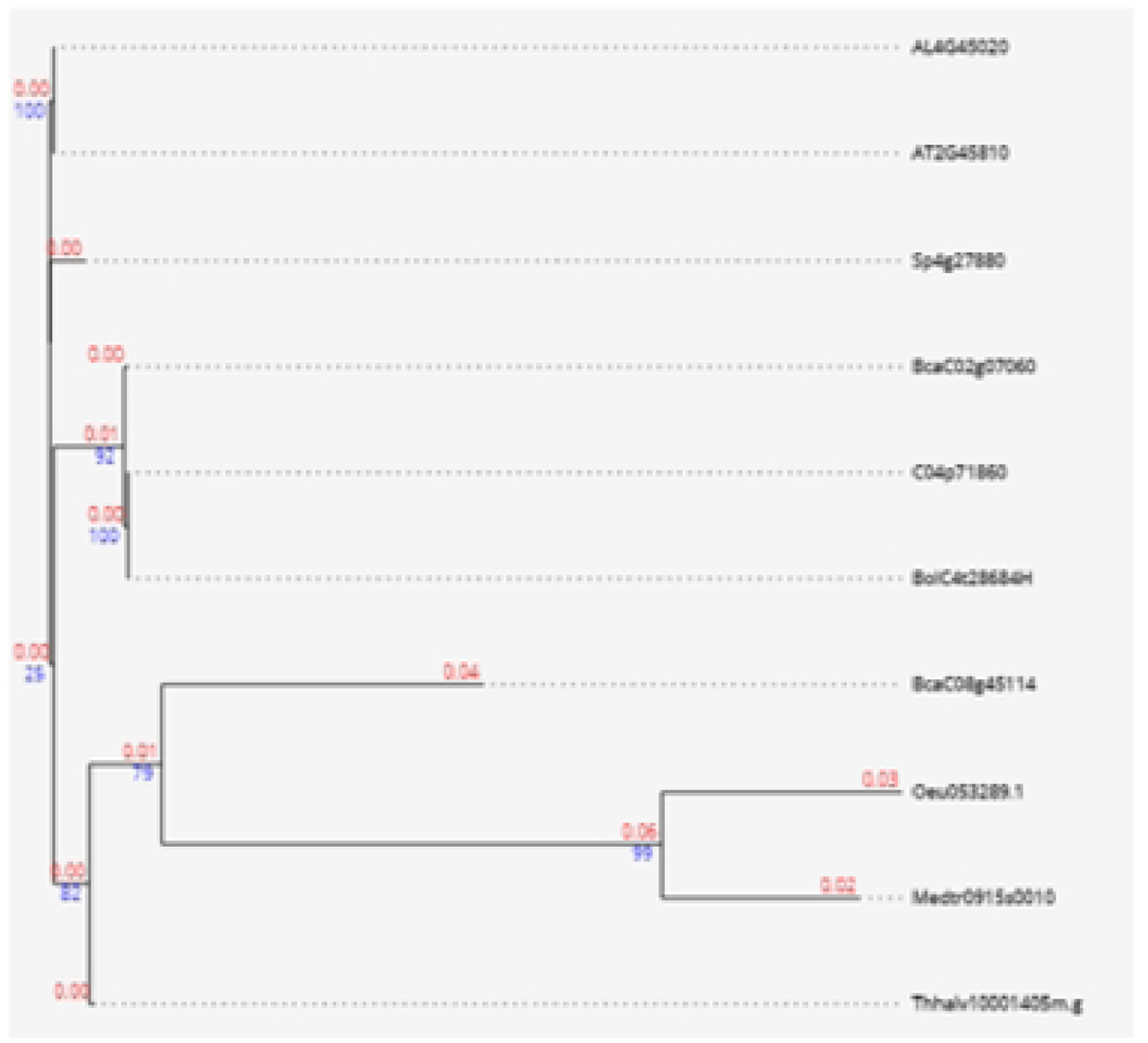
Custom-generated phylogenetic tree from PLAZA server supports the phylogenetic relationship of in-paralogs as seen in the phylogenetic tree.

### 3.3 Early developmental stages analysis

Through the computational database Arabidopsis eFP Browser (https://bar.utoronto.ca/efp/cgi-bin/efpWeb.cgi), we generated crucial development maps, embryo development, single cell analysis, DNA damage, etc., In table 2. we encompassed information on the tissue, expression value, standard deviation, and samples of gene AT2G45810 *Arabidopsis thaliana.* We found most expression level is 329.47 in imbibed seed for sample RIKEN-NAKABAYASHI2A, RIKEN-NAKABAYASHI2B.

**Table 2.**
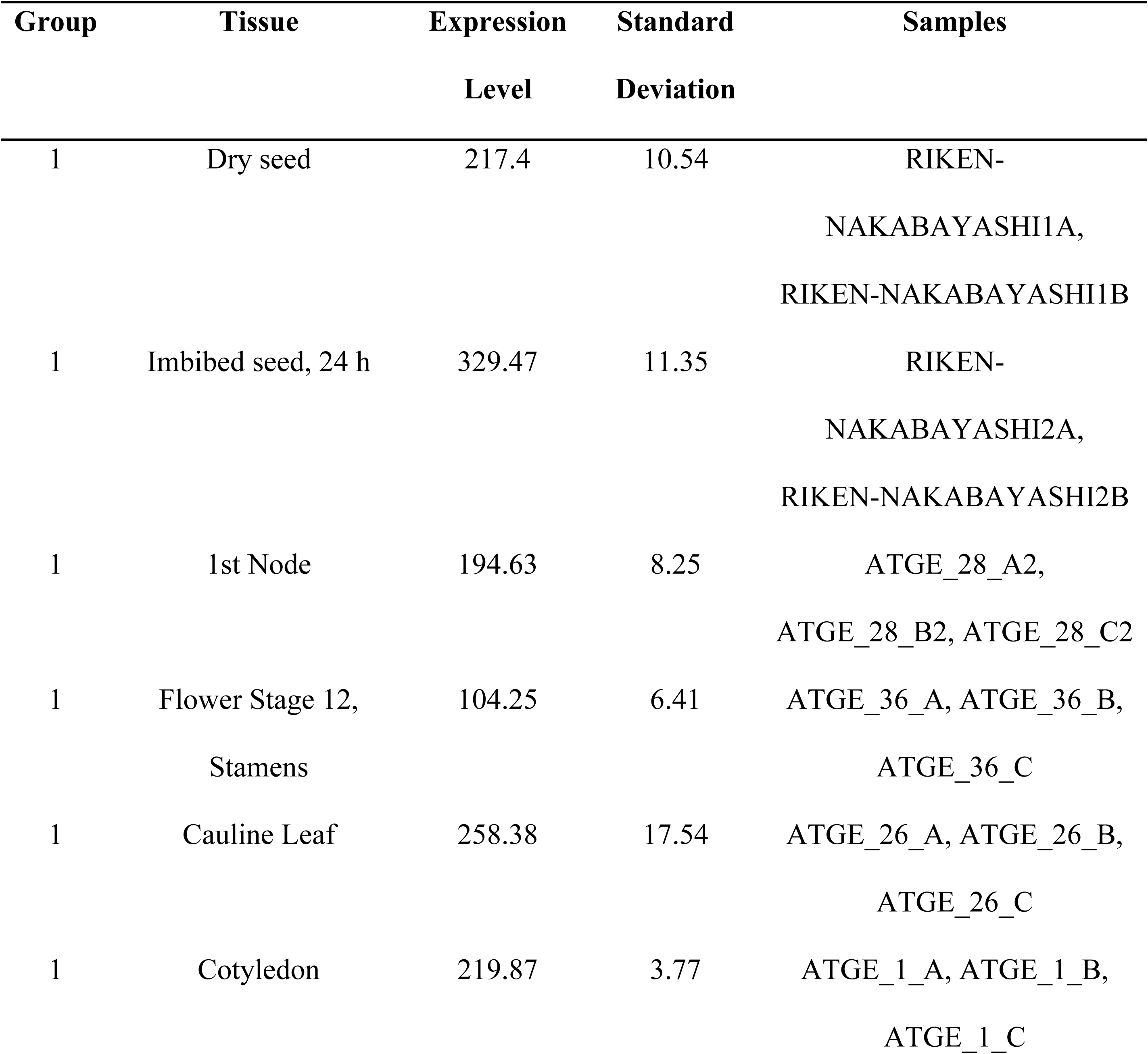
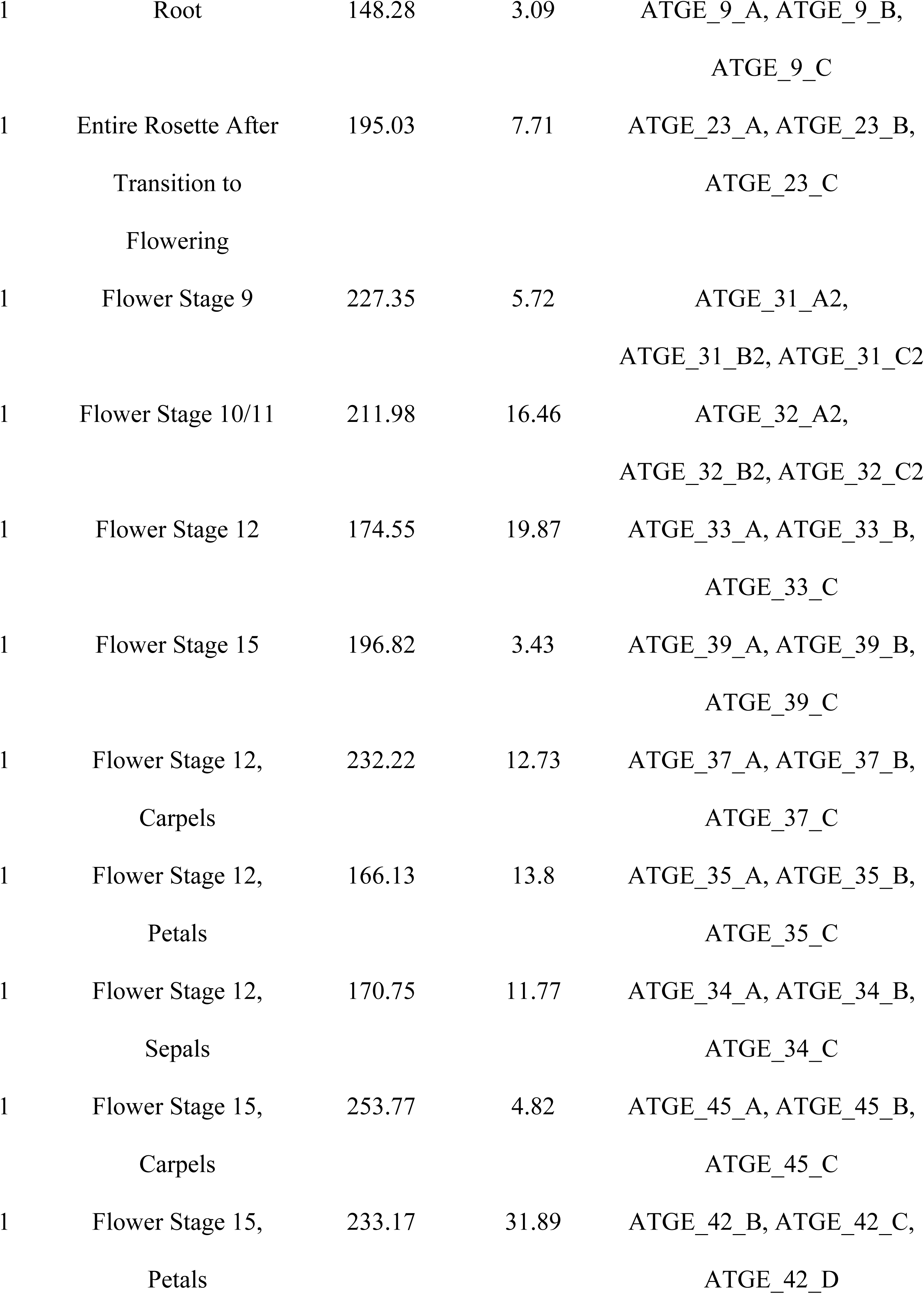
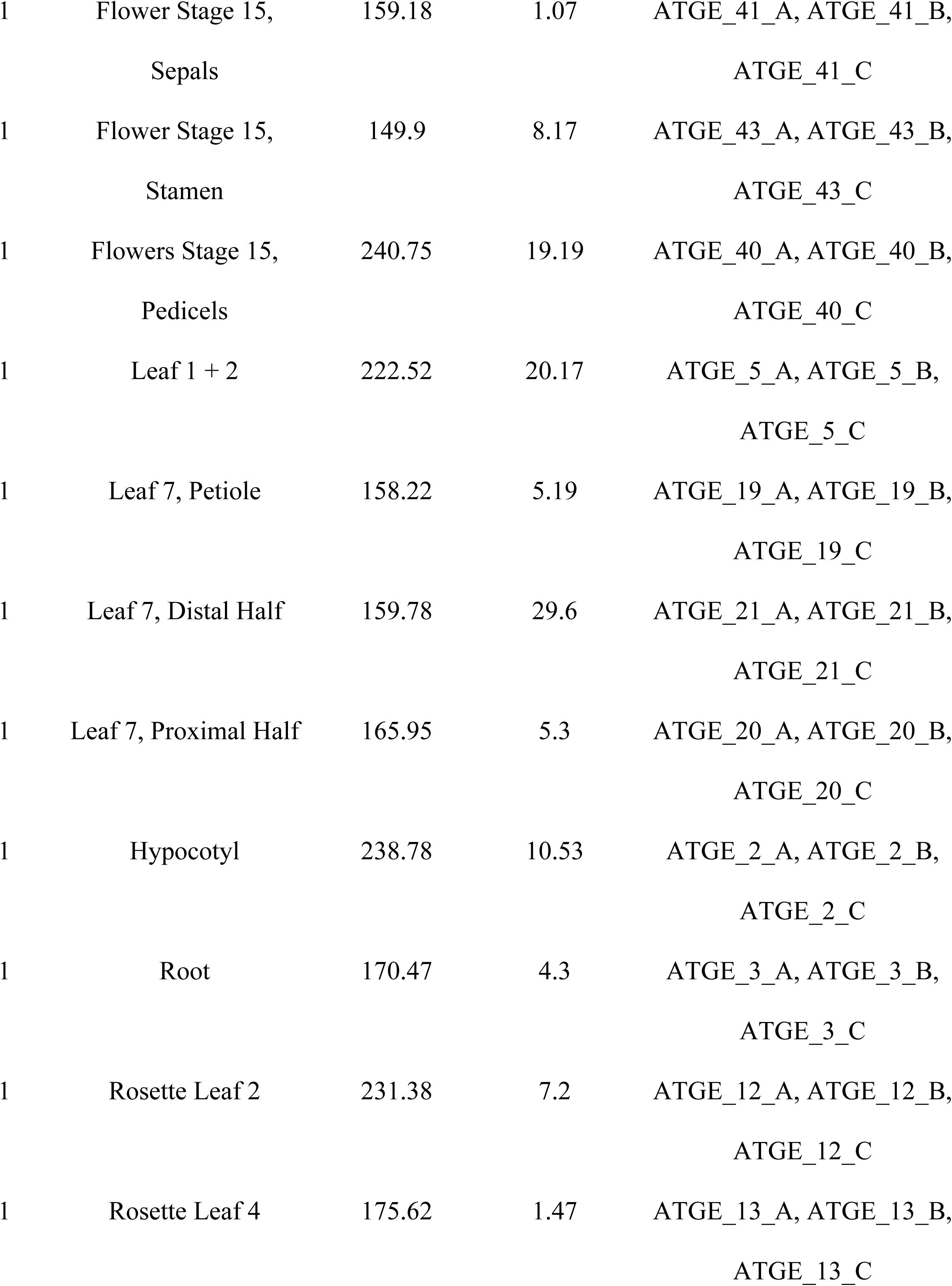
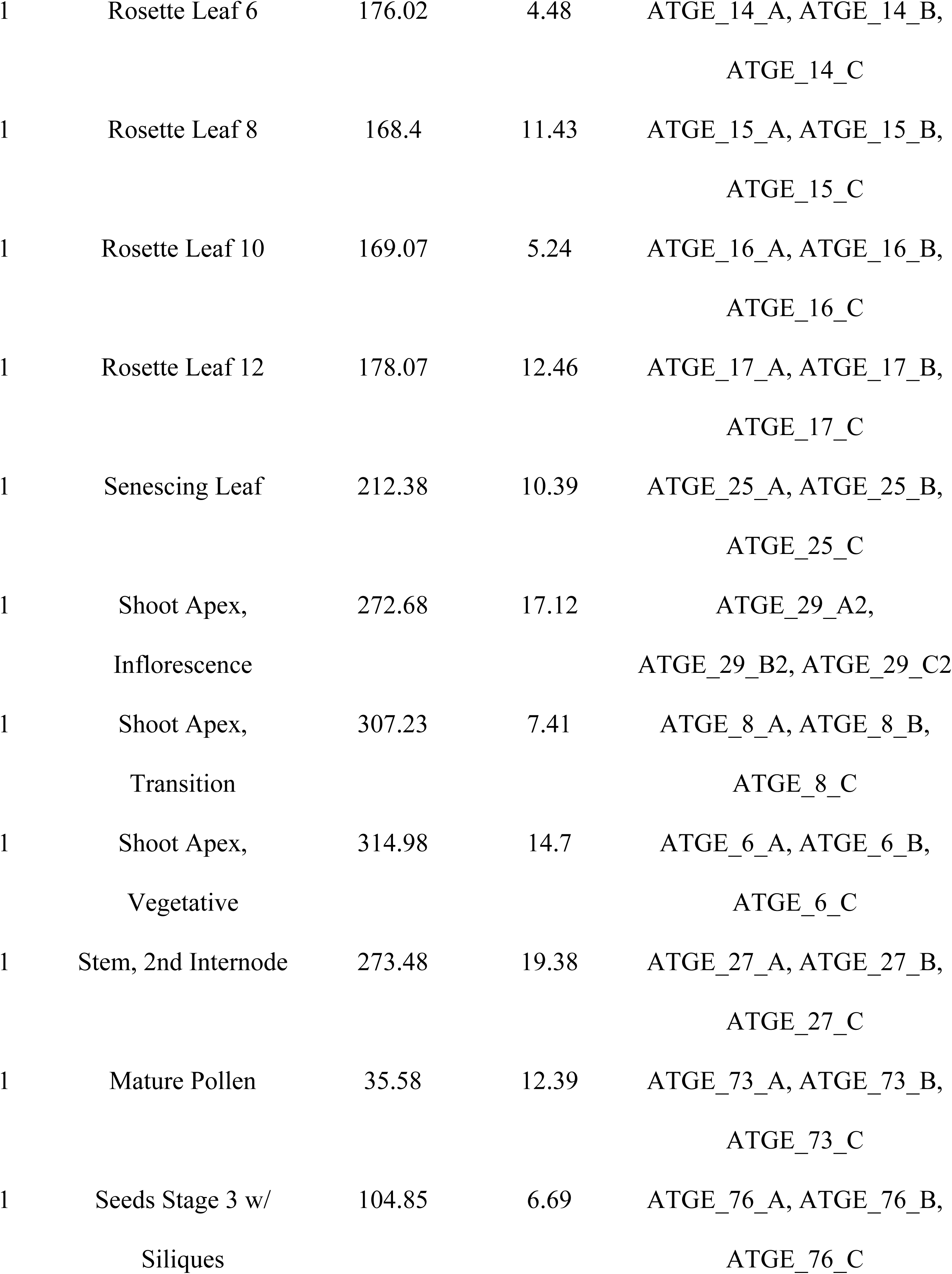
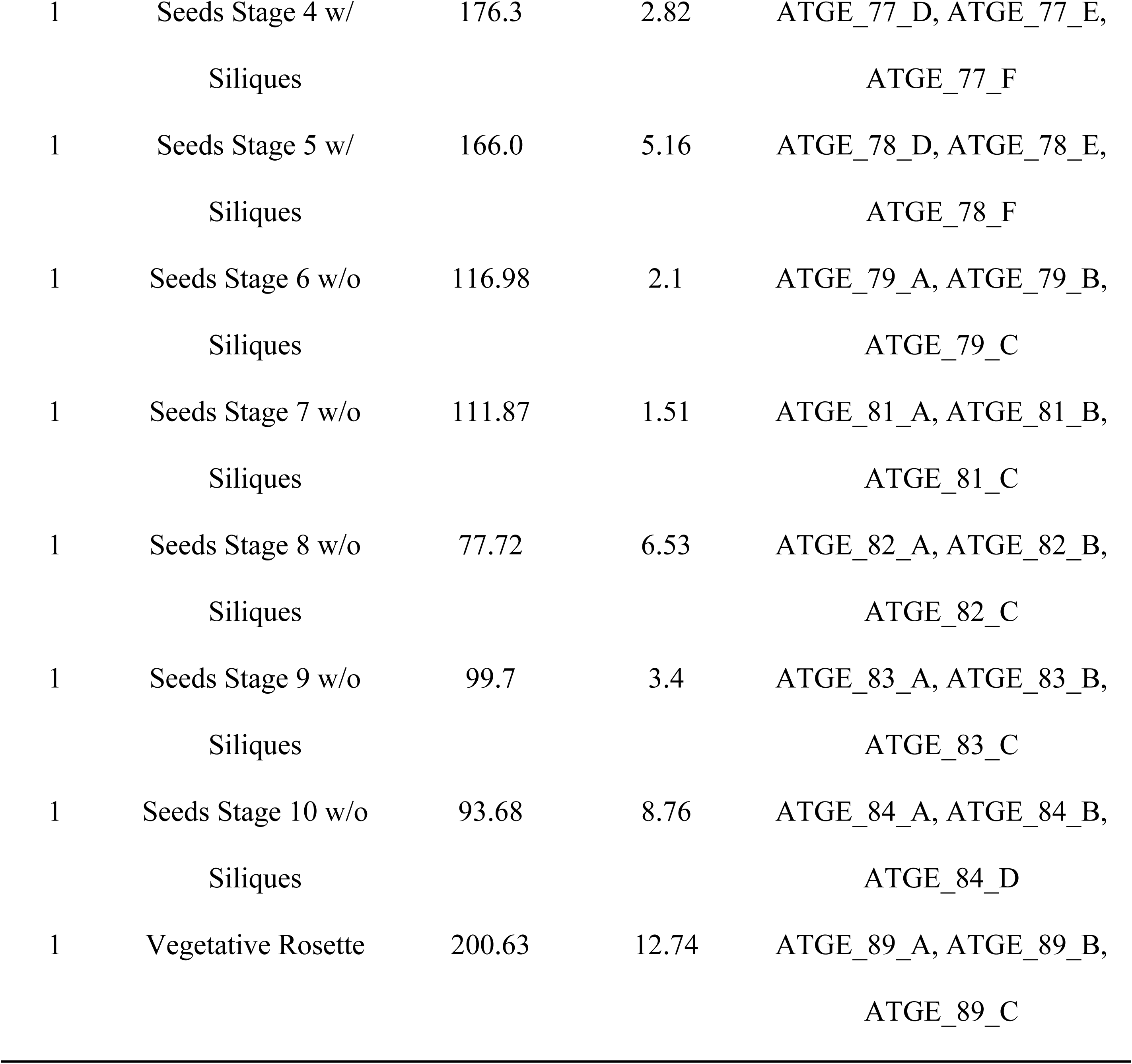
The expression value of Arabidopsis thaliana (AT2G45810): Integrated to information of the tissue, expression value, standard deviation and samples.

**Table 3.**
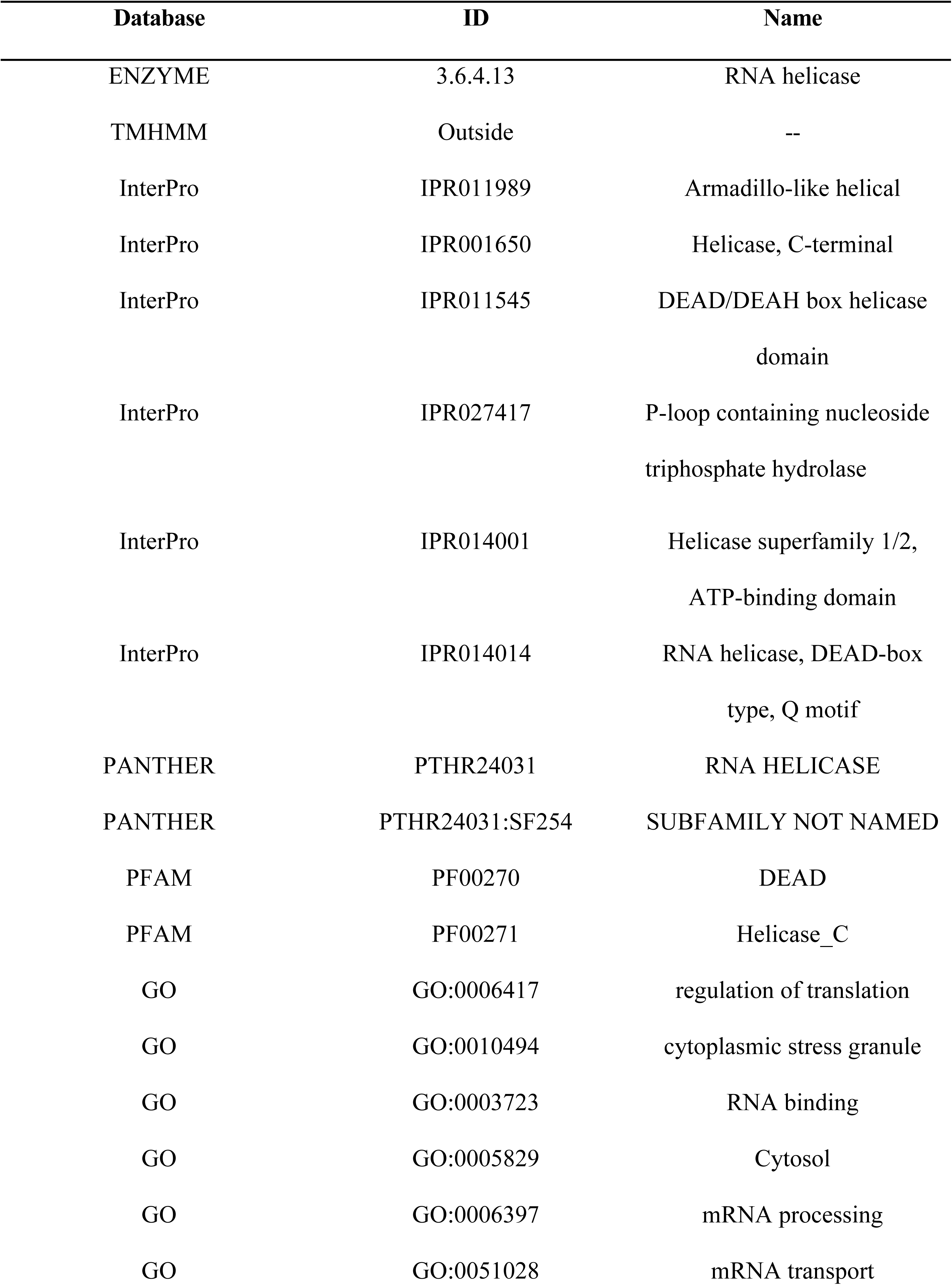
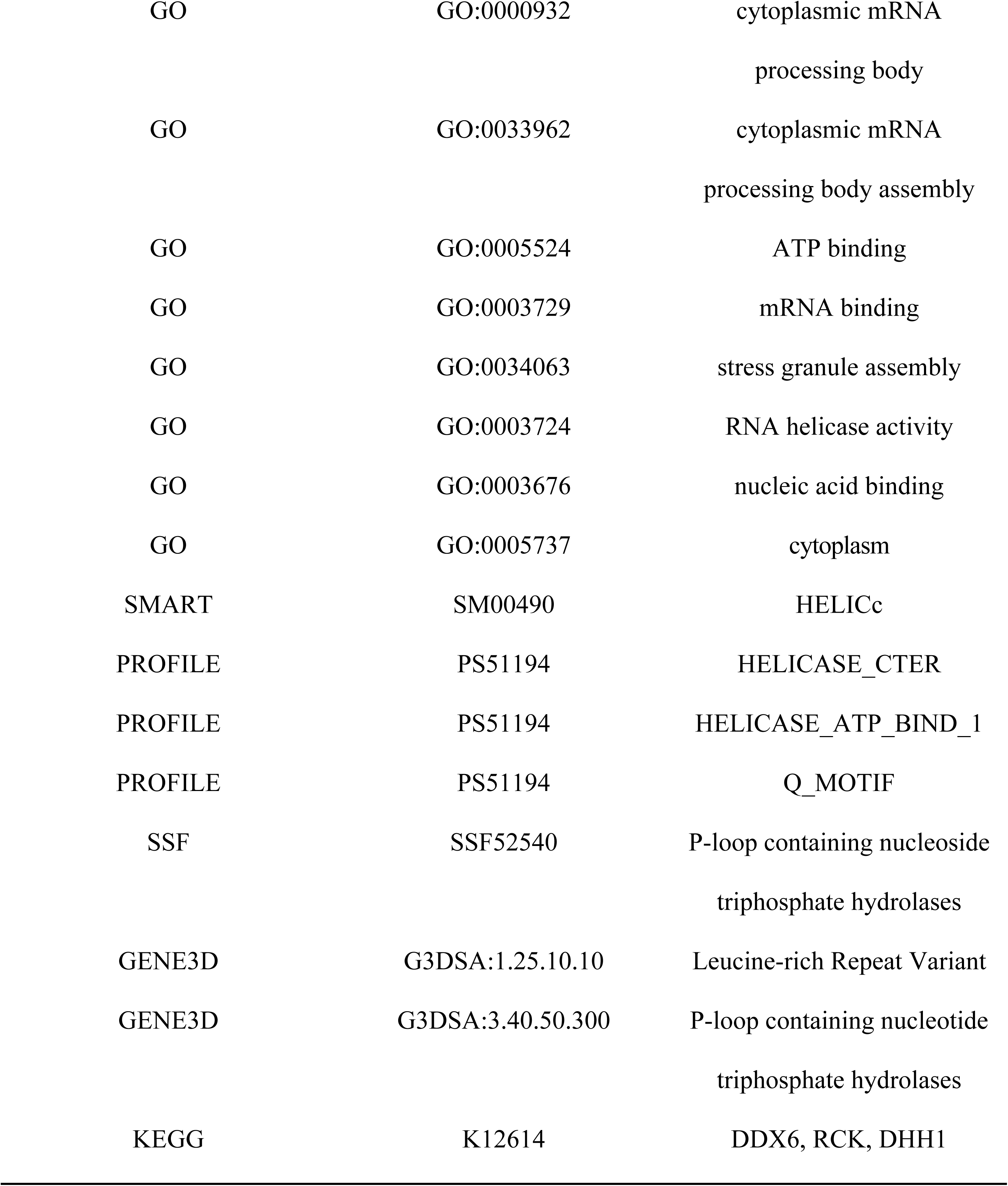
Functional annotation for Arabidopsis thaliana airport11 AT2G45810 gene.

**Fig 3.**
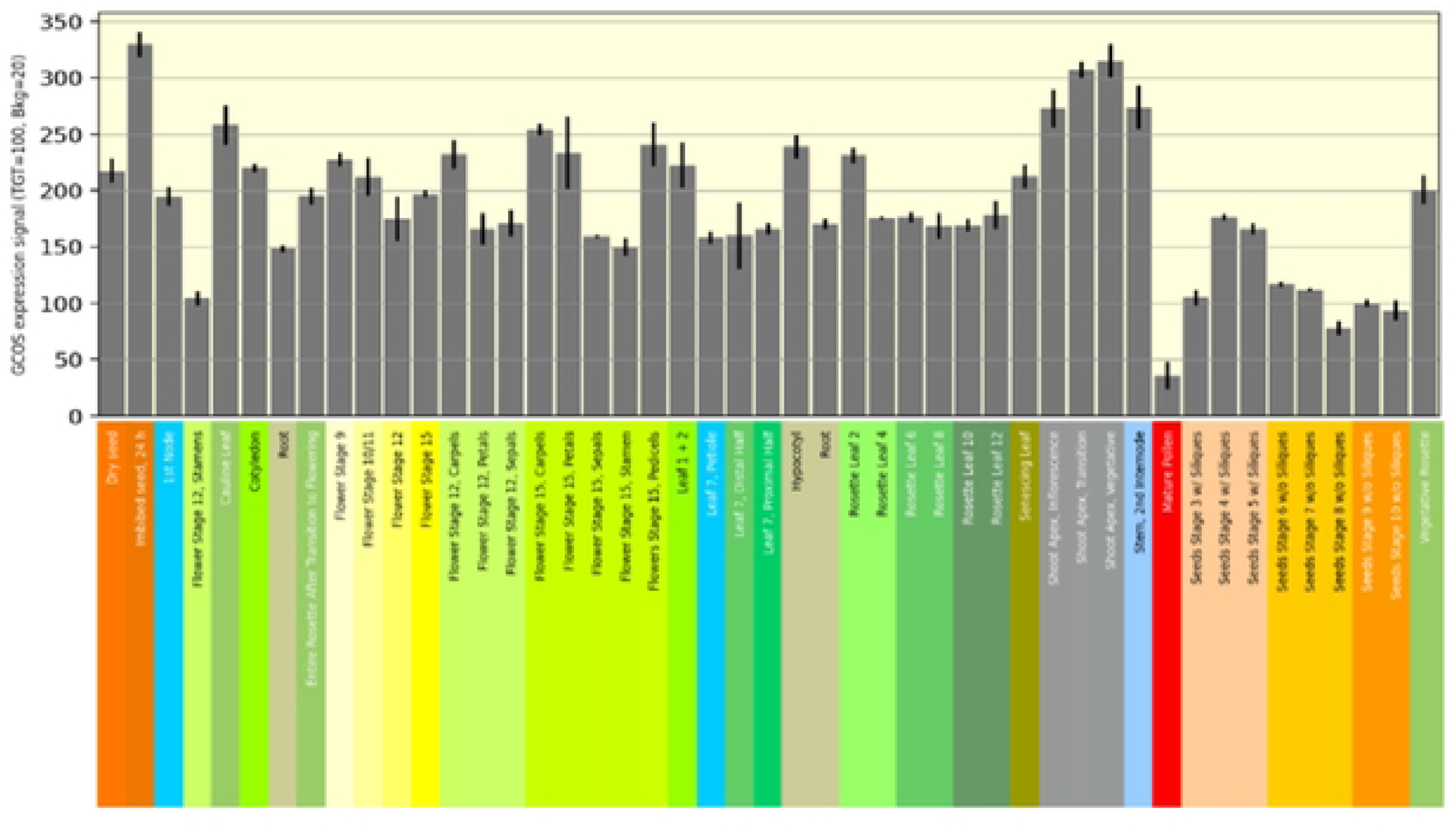
Chart of expression value (AT2G45810): GCO5 expression signal (TGT=100, Bkg=20) comparisons with several Arabidopsis thaliana physical parts.

#### Arabidopsis development map

A gene expression map may be used to examine global gene expression throughout the development of the reference plant *Arabidopsis thaliana* in samples from embryonic to senescence and across numerous organs. As a result, several transcriptional processes underpinning the development of many organ systems are always active(28,44).

#### Arabidopsis embryo

*Arabidopsis thaliana* plant have a unique and special embryo development stages that described in figure 4(18). Col-0 seedlings were cultivated in an environmentally controlled growing room at a temperature of 20-22 °C with a 16-hour light/8-hour dark cycle. For quantification of all mRNA seq datasets, the pseudo-aligner Kallisto (v0.44 0) was utilized (45). Before measurement, the essential adapter sequences for each mRNA-seq collection were trimmed from the FASTQ files using Cutadapt with a match length of at least five nucleotides(46).

**Fig 4.**
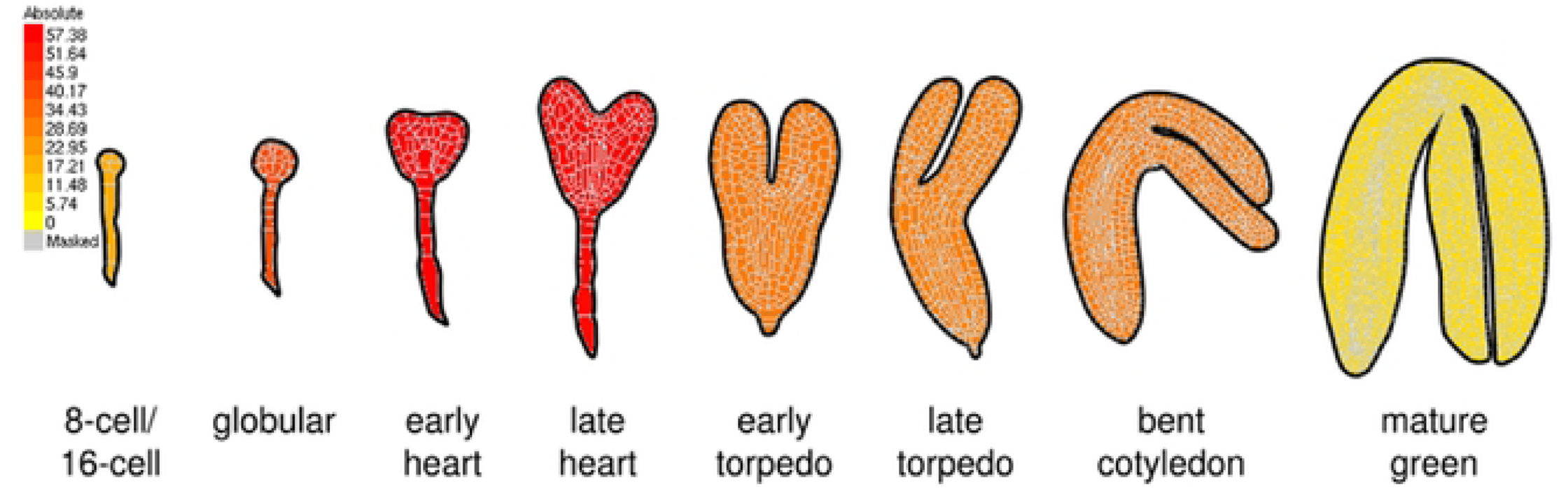
Arabidopsis embryo developmental stages.

#### Single-cell analysis

Using the default settings, raw reads were demultiplexed and mapped to the TAIR1O reference genome by the 1OX Genomics Cell Ranger pipeline (v2.1,1). Unless otherwise specified, all downstream single-cell analyses were carried out using Seurat (v2.3.3)(47)(48). In summary, unique molecular IDs were tallied for each gene and each cell barcode (filtered by Cell Ranger) to create digital expression maps. A gene was deemed expressed if it was expressed in more than three cells, and each cell had to contain at least 200 expressed genes.

#### DNA damages

ONA Double Strand Breaks are formed as a result of gamma irradiation (V-IR) (OSBs; marked bV stars). When these breaks are recognized, the ATAXIA-telangiectasla mutated (ATM)-dependent phosphorylation of suppressor of gamma 1 (SOG-l), a master regulator of the ONA damage response, occurs (filled circles).(49) SOG-l is a MAC domain transcription factor found through a DNA damage suppressor screening that has been functionally correlated with the human tumor suppressor gene, p53(50)(51)(52).

### 3.4 Expression comparison via scRNA viewer

We predicted several parameters such as time group, region, cell type, developmental stage, expression and time through the scRNA viewer computation tool with comparison their expression level.

**Fig 5.**
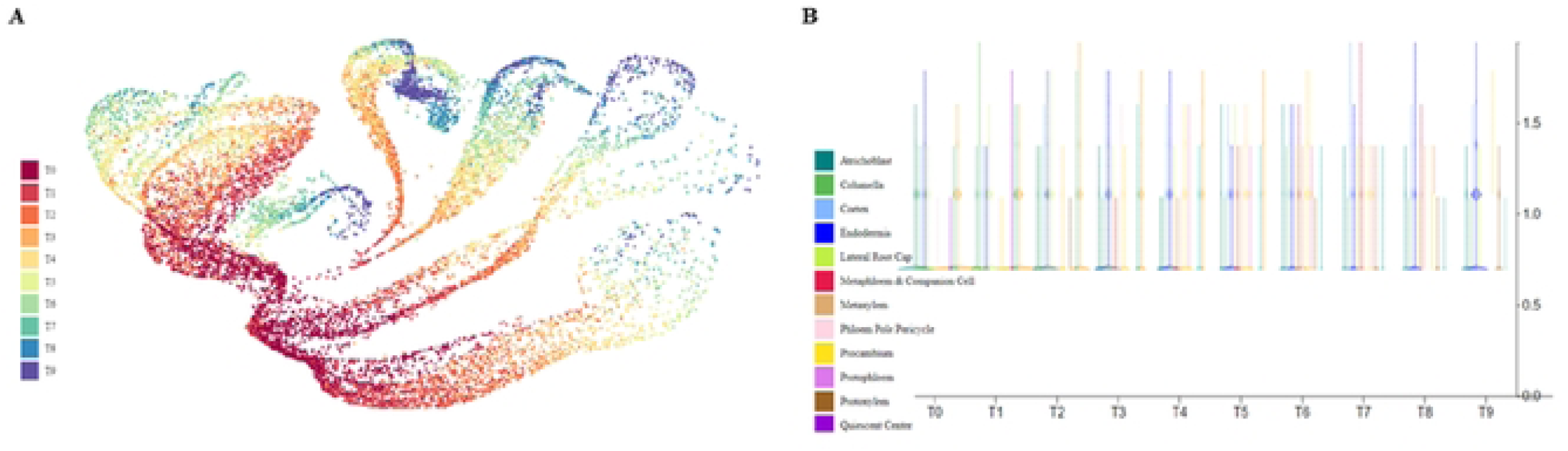
Arabi Atlas Data Expression for gene AT2G45810 from scRNA viewer. A. (Time Group) B. (Region)

**Fig 6.**
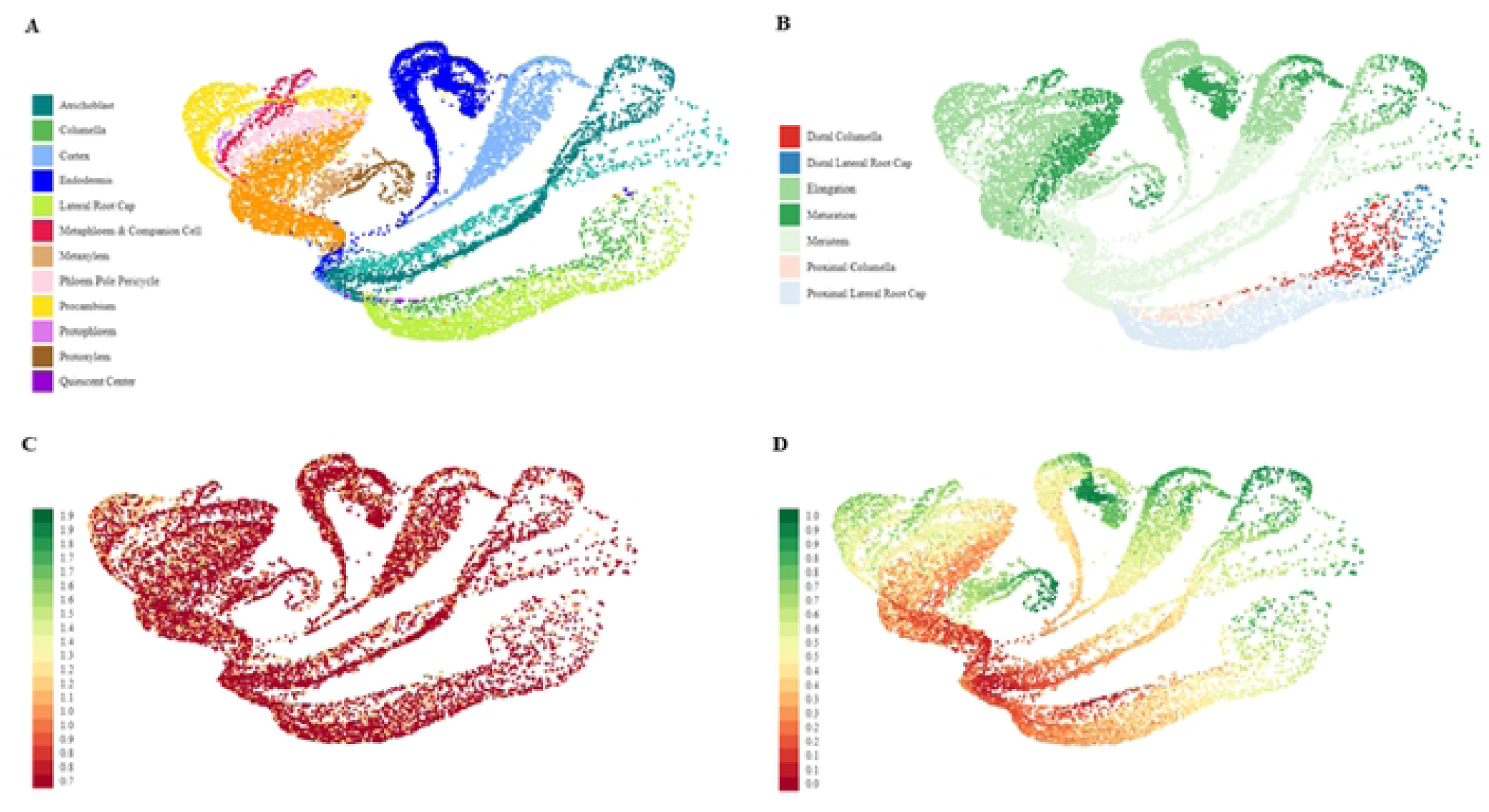
Arabi Atlas Data Expression for gene AT2G45810 from scRNA viewer. A. (Cell Type) B. (Development Stage) C. (Expression) D. (Time)

### 3.5 Functional Annotations

We analyzed the functional annotations of gene AT2G45810 through several online servers including ENZYME (https://enzyme.expasy.org/)(53), Interpro (https://www.ebi.ac.uk/interpro/)(54), PANTHER (https://www.pantherdb.org/)(55)(56), Pfam (https://pfam.xfam.org/), (57) QuickGo (https://www.ebi.ac.uk/QuickGO/)(58), SMART (https://smart.embl.de/)(59)(60), PROSITE (https://prosite.expasy.org/)(61), SUPERFAMILY (https://supfam.org/SUPERFAMILY/index.html)(62), CATH (https://www.cathdb.info/)(63), and KEGG (https://www.genome.jp/kegg/kegg2.html)(64).

**Fig 7.**
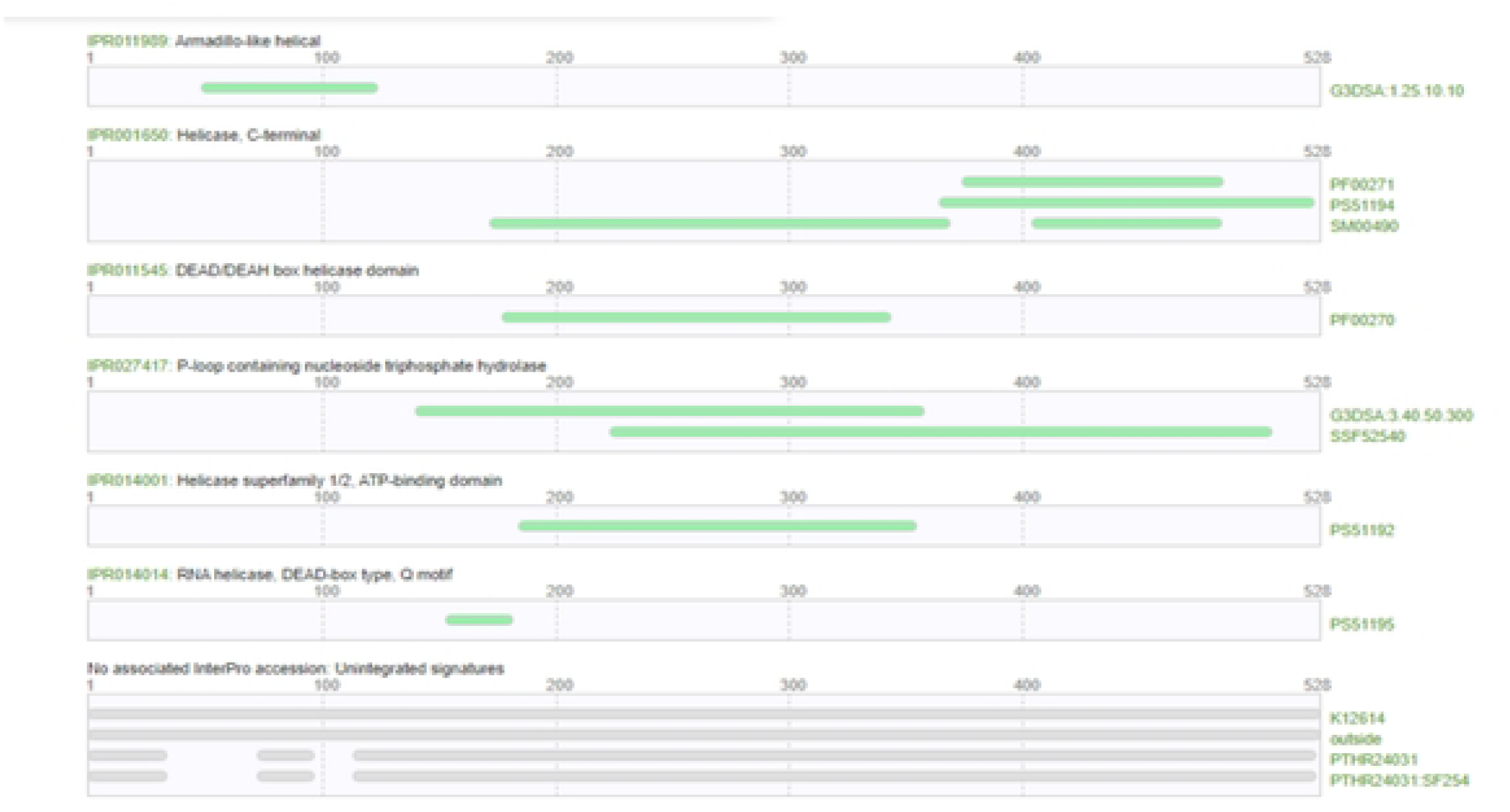
Functional annotation for Arabidopsis thaliana airport11 AT2G45810 gene.

### 3.6 Protein homologs

*A. thaliana* TAIR10 (transcript: AT2G45810.1) DEA(D/H)-box RNA helicase family protein have 97.5% similarity with *A. lyrata* v2.1 (transcript: AL4G45020.t1) and define as (1 of 3) K12614 - ATP-dependent RNA helicase DDX6/DHH1 (DDj). The comprehensive information at protein homologs of *Arabidopsis thaliana* (AT2G45810) enlisted in table 4.

**Table 4.**
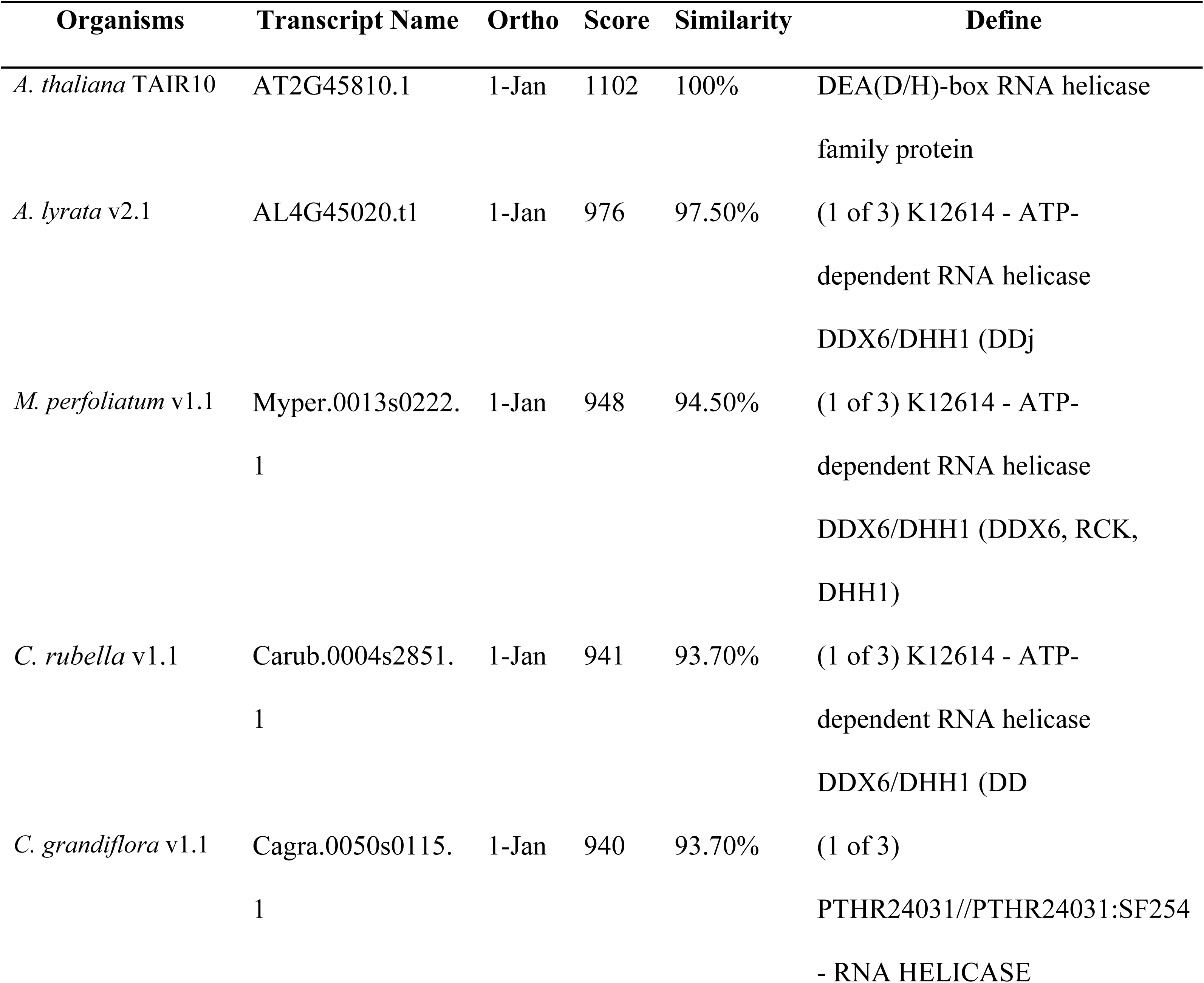
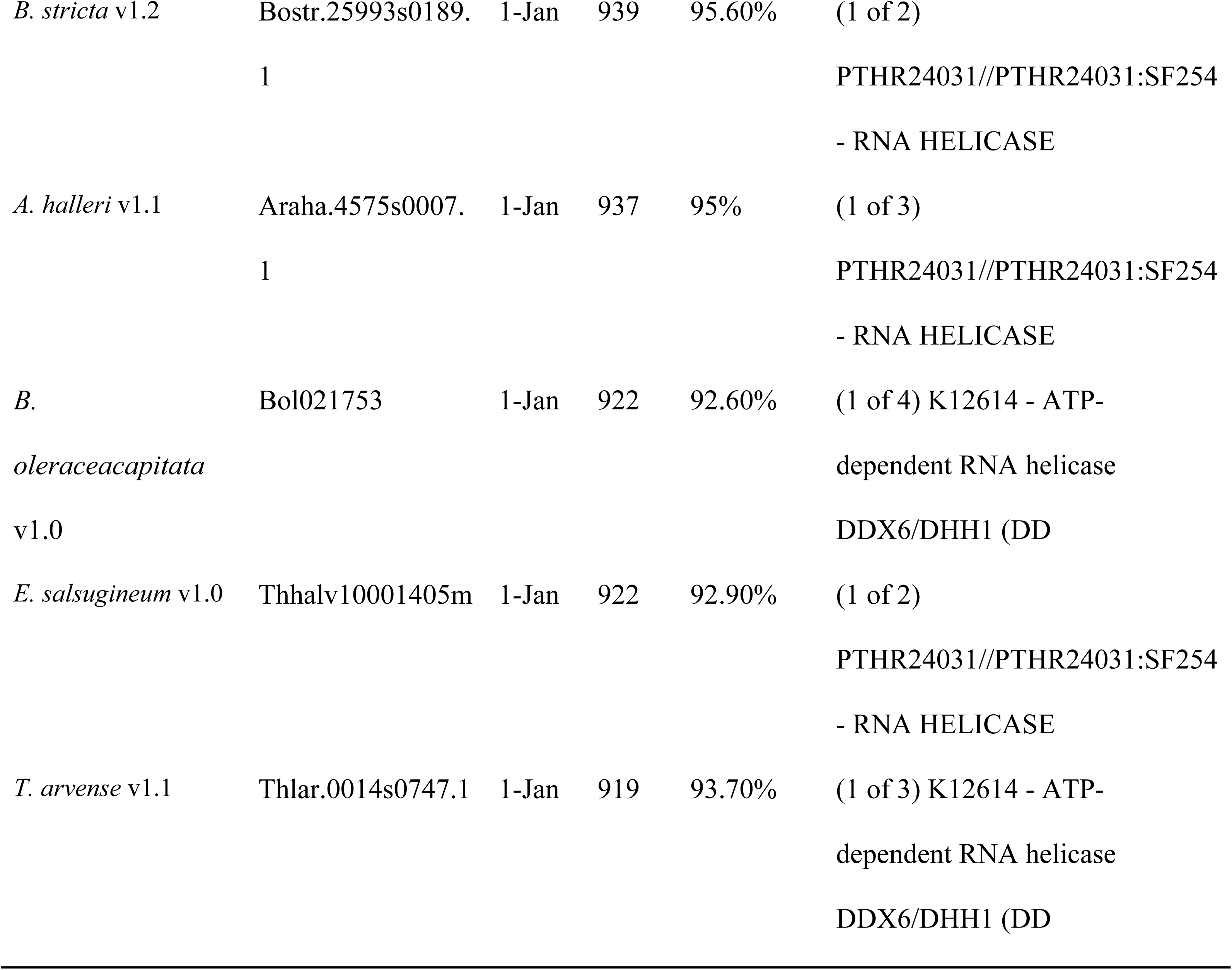
Protein homologs of Arabidopsis thaliana (AT2G45810).

### 3.7 PlantFAMs bio sequence analysis

*Arabidopsis thaliana* airport11 AT2G45810 gene associated PlantFAMs bio sequence analysis using profile hidden Markov models using HMMER. Node Viridiplantae protein family DEA(D/H)-box RNA helicase (Family ID: 122825990) showed score 1187.9 with family size strain. DEAD-box ATP-dependent RNA helicase, putative, expressed, K00753 - glycoprotein 3-alpha-L-fucosyltransferase (E2.4.1.214), ATP-dependent RNA helicase, and K13179 - ATP-dependent RNA helicase DDX18/HAS1 [EC:3.6.4.13] (DDX18, HAS also showed significant score. Further information described in the table 5 at associated PlantFAMs bio sequence analysis using profile hidden Markov models.

**Table 5.** Associated PlantFAMs bio sequence analysis using profile hidden Markov models using HMMER.

| Node | Name | Family ID | Score | Significance | Family size |
| --- | --- | --- | --- | --- | --- |
| Viridiplantae | DEA(D/H)-box RNA helicase<br>family protein | 122825990 | 1187.9 | 0 | 8 |
| Viridiplantae | DEA(D/H)-box RNA helicase<br>family protein | 122825968 | 1025 | 0 | 8 |
| Viridiplantae | DEAD-box ATP-dependent RNA<br>helicase, putative, expressed | 123424669 | 979.8 | 2.3e-287 | 136 |
| Viridiplantae | K00753 - glycoprotein 3-alpha-L-fucosyltransferase (E2.4.1.214) | 123084736 | 896.7 | 3.5e-262 | 2 |
| Viridiplantae | K12614 - ATP-dependent RNA<br>helicase DDX6/DHH1 (DDX6,<br>RCK, DHH1) | 123281937 | 874 | 1.8e-255 | 2 |
| Viridiplantae | PTHR24031//PTHR24031:SF254<br>- RNA helicase // subfamily not<br>named | 123274745 | 759.8 | 4.7e-221 | 3 |
| Viridiplantae | ATP-dependent RNA helicase | 123095611 | 350.2 | 1.4e-96 | 2 |
| Viridiplantae | DEAD box RNA helicase family<br>protein | 123429150 | 338.3 | 9.8e-93 | 527 |
| Viridiplantae | DEAD-box ATP-dependent RNA<br>helicase, putative, expressed | 123424878 | 325.9 | 3.6e-89 | 178 |
| Viridiplantae | K13179 - ATP-dependent RNA<br>helicase DDX18/HAS1<br>[EC:3.6.4.13] (DDX18, HAS1) | 123429061 | 311.1 | 6e-85 | 162 |

**Fig 8.**
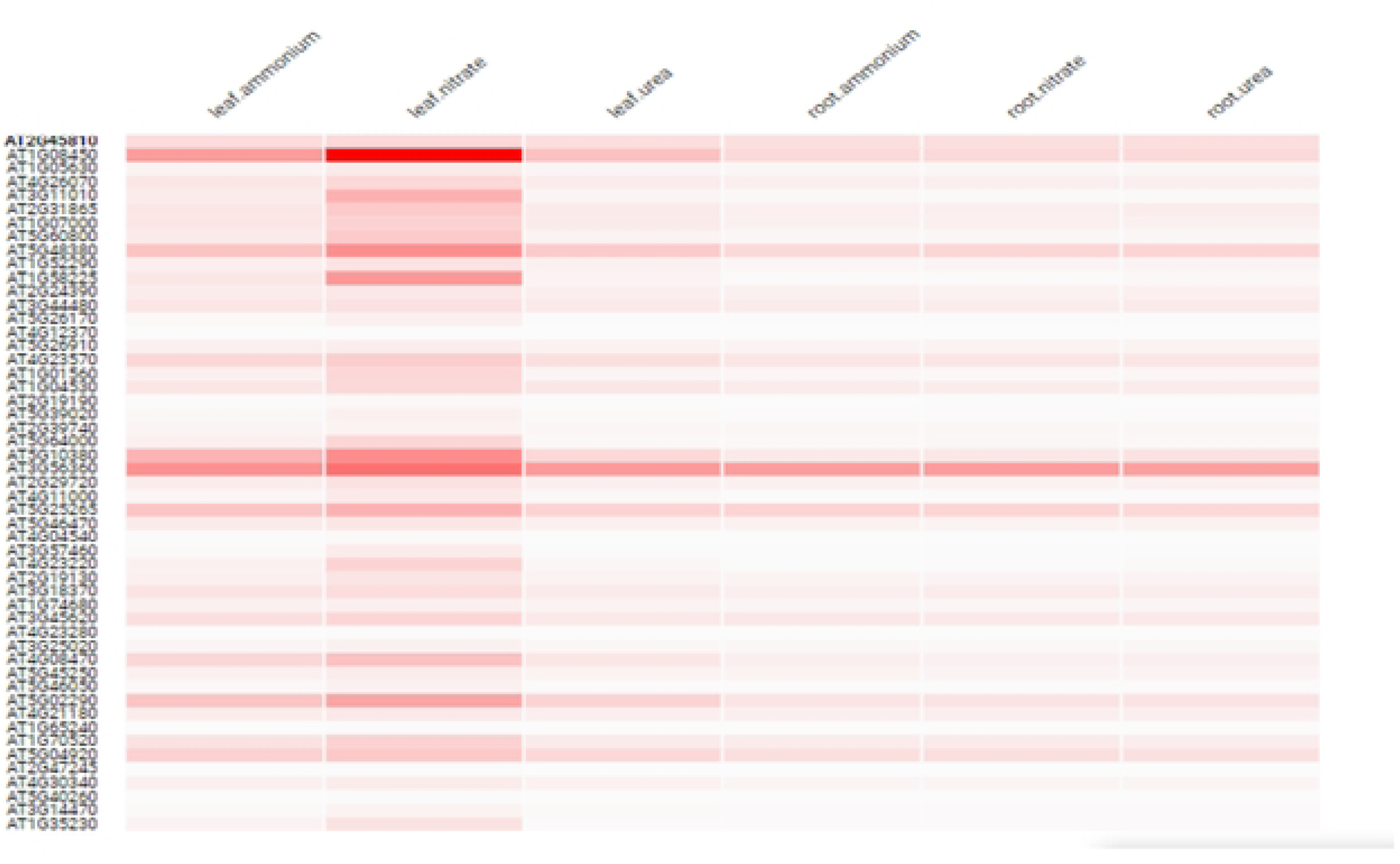
Arabidopsis thaliana gene AT2G45810 expression graph integrated to most co-expressed genes, GeneAtlas v2 FPKM experiment group.

### 3.8 Co-expression analysis

We used the BAR utoronto’s Expression Angler for the determined and analysis of the Co-expression profile for the specific gene At2g45810. The top 50 co-expressed genes with At2g45810 in the Developmental Map view were downloaded and were compared to the top 50 co-expressed with a “Custom Expression Pattern” as defined by At2g45810 in the Tissue Specific view.

**Fig 9.**
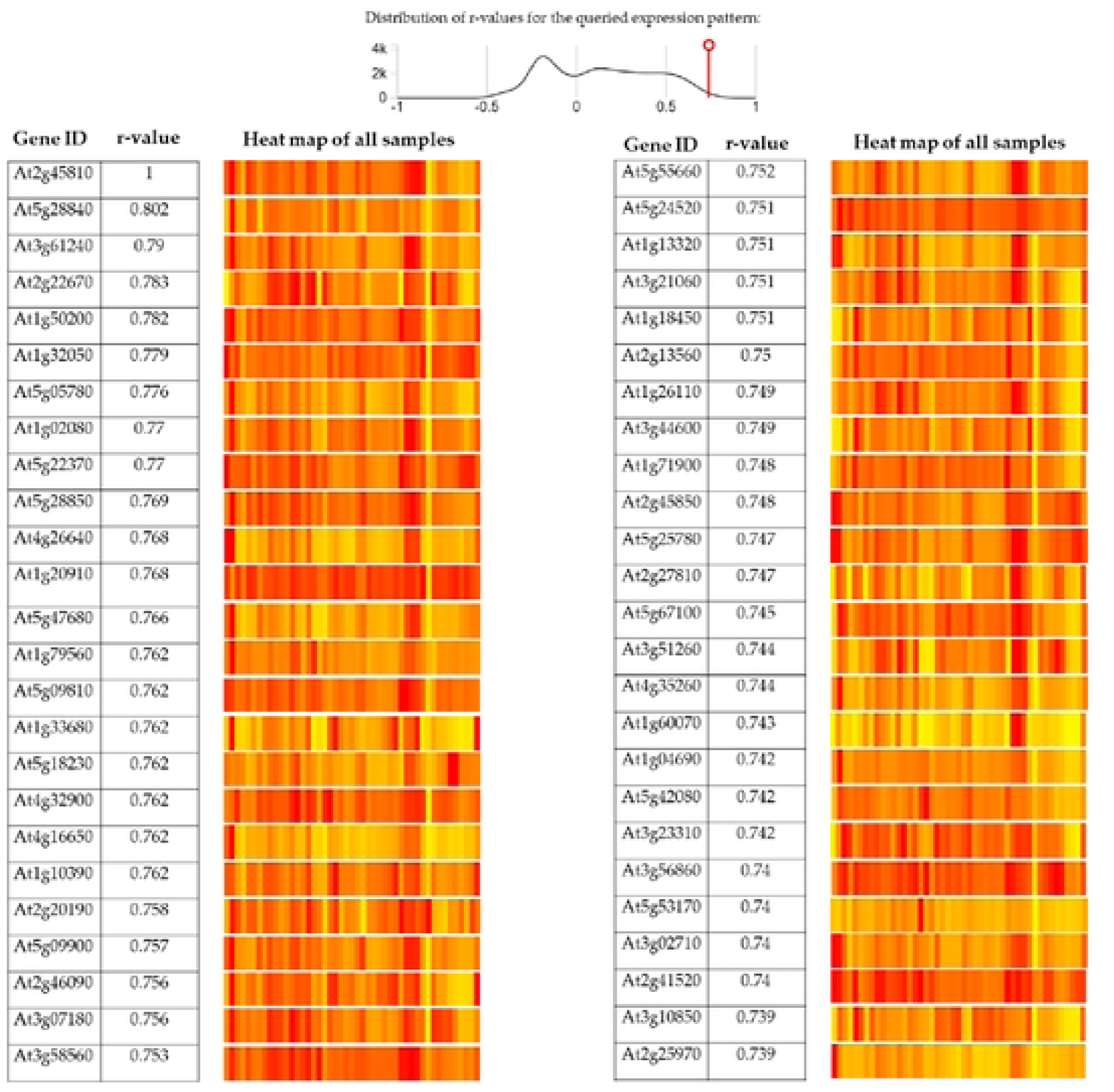
Defining a Custom Expression Pattern that matches AT2G45810 in the Tissue Specific view.

### 3.9 Promoter analysis

The Bio-Analytic Resource (BAR) widely used and scientifical computational Cistome tools shows that the 26 motifs. AGIs without any (significantly enriched) cis-elements: AT2G45810 AT5G28840 AT3G61240 AT2G22670 AT1G50200 AT1G32050 AT5G05780 AT5G22370 AT1G02080 AT5G28850 AT1G79560 AT5G09810 AT4G32900 AT5G09900 AT2G46090 AT3G07180 AT3G58560 AT1G13320 AT1G18450 AT2G13560 AT1G26110 AT1G71900 AT5G25780 AT4G35260 AT3G23310 AT5G42080 AT3G56860 AT2G41520 AT3G02710 AT2G25970 AT5G38470. Functional depth cutoff was between 0 and 1 (0.75). Promoter identification and analysis showed in fig 10.

**Fig 10.**
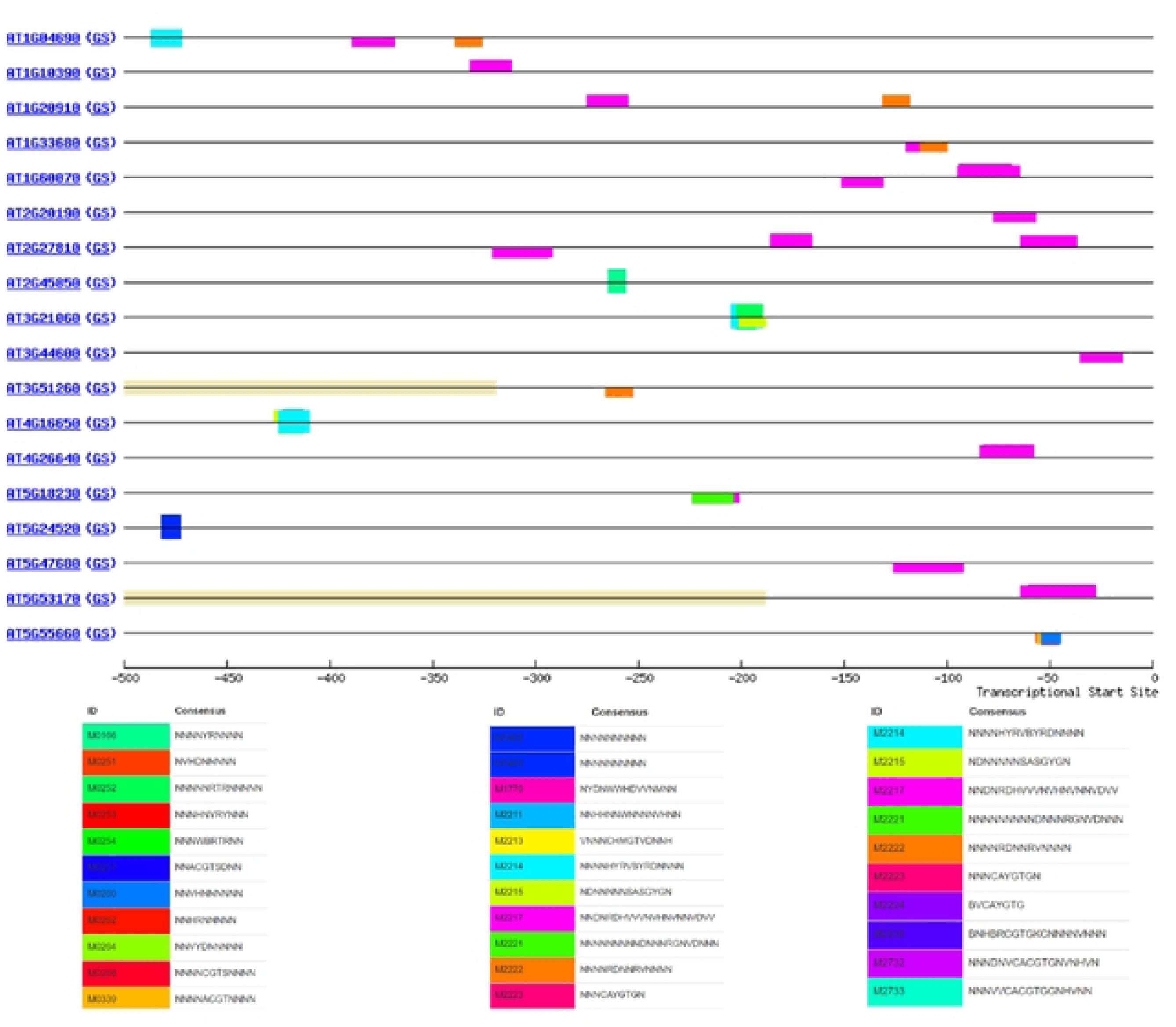
The motif enriched in the promoters of genes co-expressed with the same “tissue specific” pattern of expression in pollen and guard cells as AT2G45810.

### 3.11 Functional classification and pathway visualization

The AgriGO analysis showed and predicted that the 50 co expression genes with the same tissue specific expression pattern as AT2G45810. We have analyzed significant levels of the alliase generated by co expression top 50 genes. We also investigated the AGI’s term name, term ID, Padj, −log10(Padj) (0 to ≤ 16) information on the basis of GO: MF, GO: BP, GO: CC, KEGG, and WP. Pathway visualization and functional categorization was showed in the fig 11 and fig 12.

**Fig 11.**
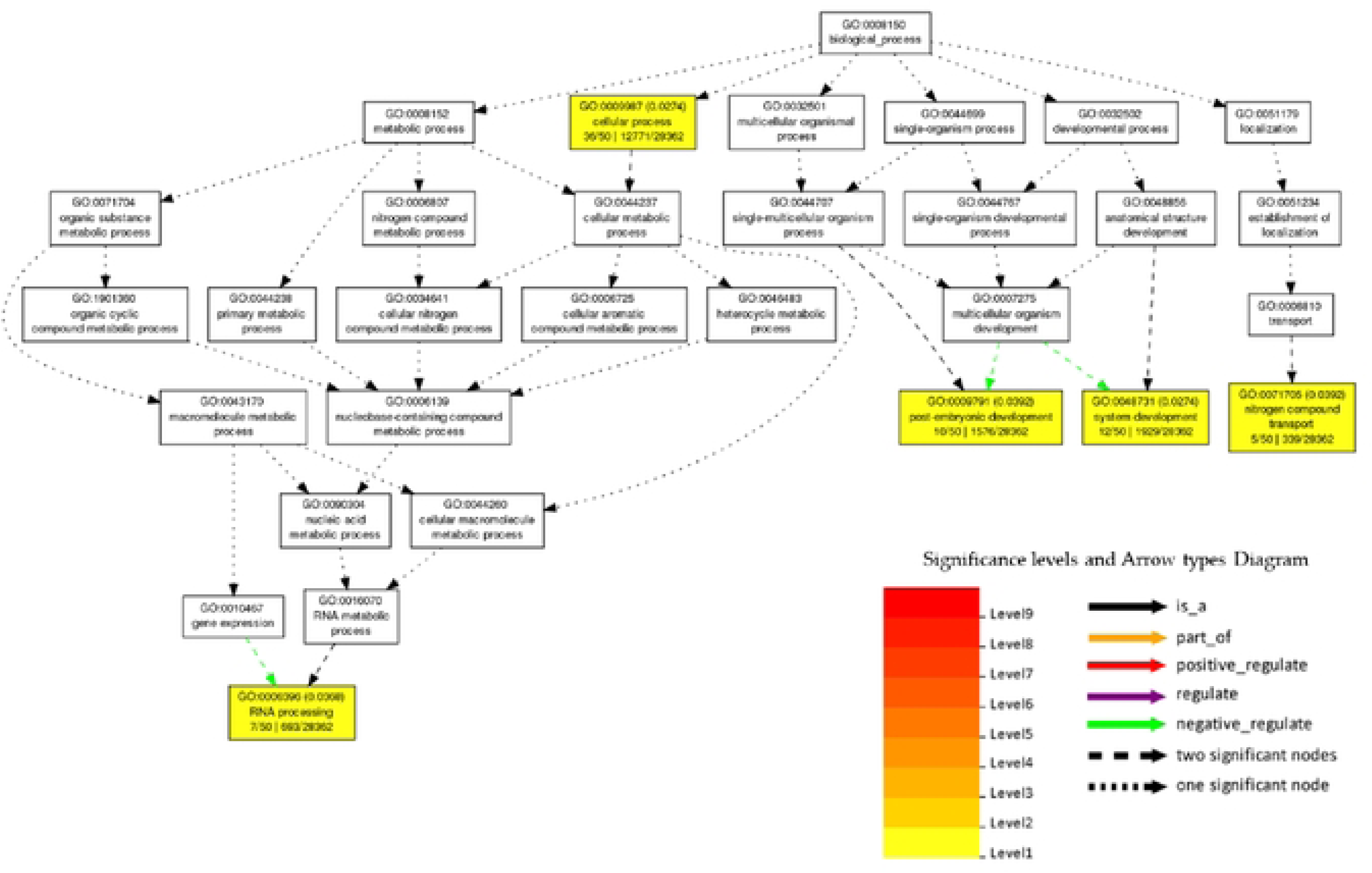
GO008150 Biological processes significant levels and arrow types diagram.

**Fig 12.**
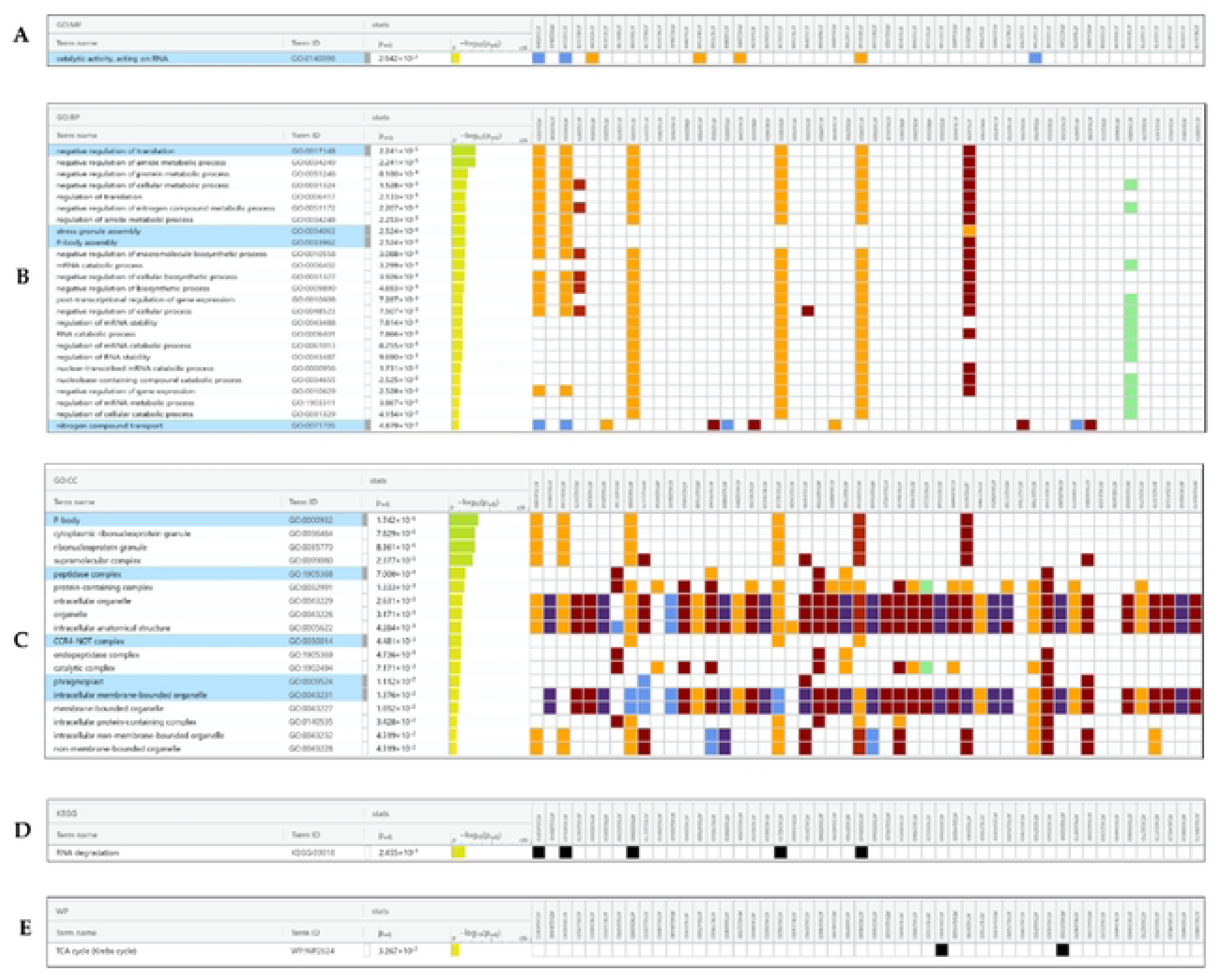
GO: MF, GO: BP, GO: CC, KEGG, and WP graph generated by AgriGO server.

### 3.12 Exploring cellular interaction

Through the computation server Arabidopsis Interactions Viewer 2 (AIV-2) we have examined cellular interaction as the protein-protein interactors of AT2G45810 and its interactors, AT5G28840 and AT3G61240, with expression levels overlaid in mature pollen. As well as predicted and analyzed protein-protein interactors of AT2G45810 and its interactors, AT5G28840 and AT3G61240, with expression levels overlaid in guard cells plus 100 um ABA.

**Fig 13.**
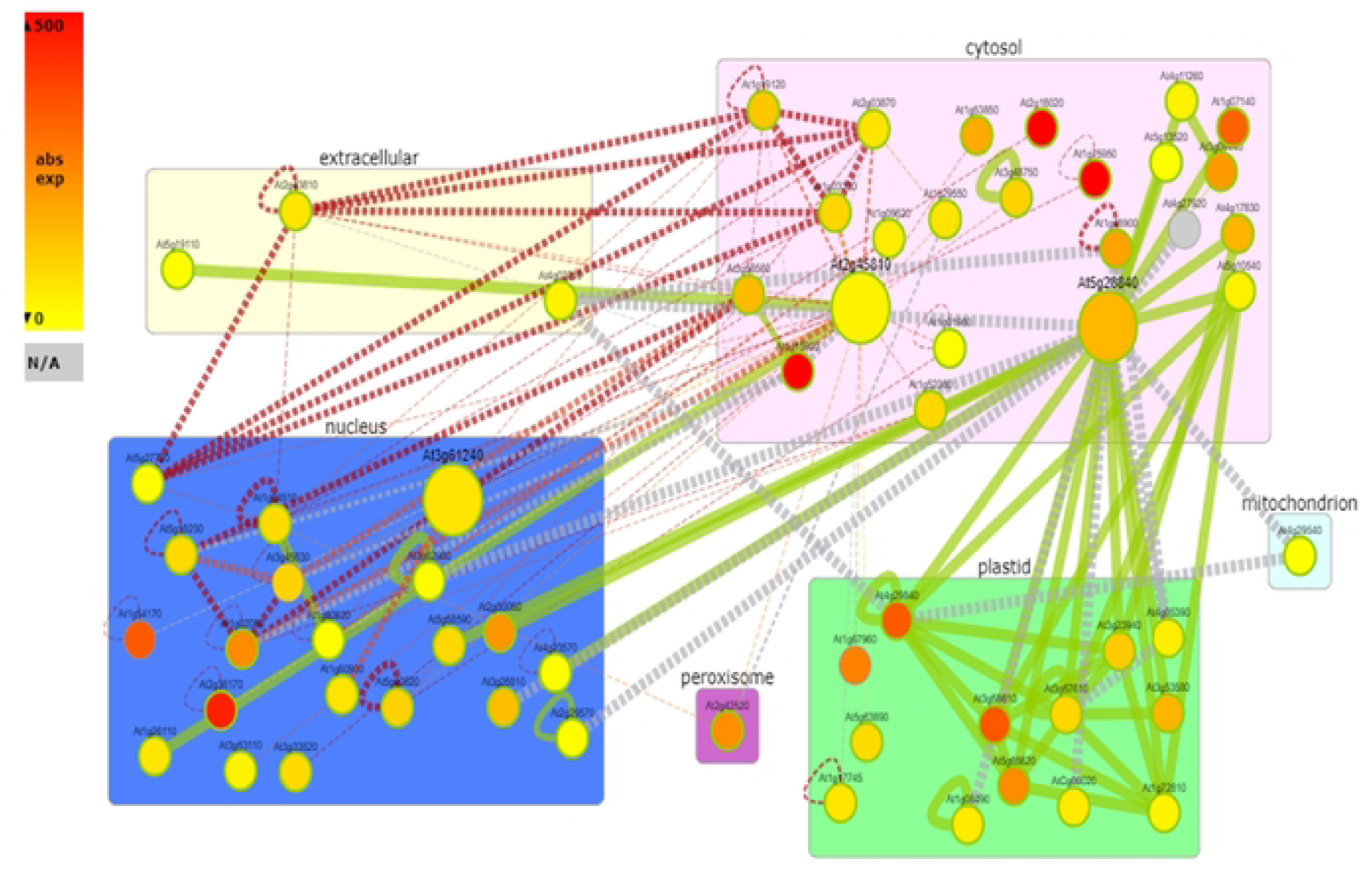
Protein-protein interactors of AT2G45810 and its interactors, AT5G28840 and AT3G61240, with expression levels overlaid in mature pollen.

**Fig 14.**
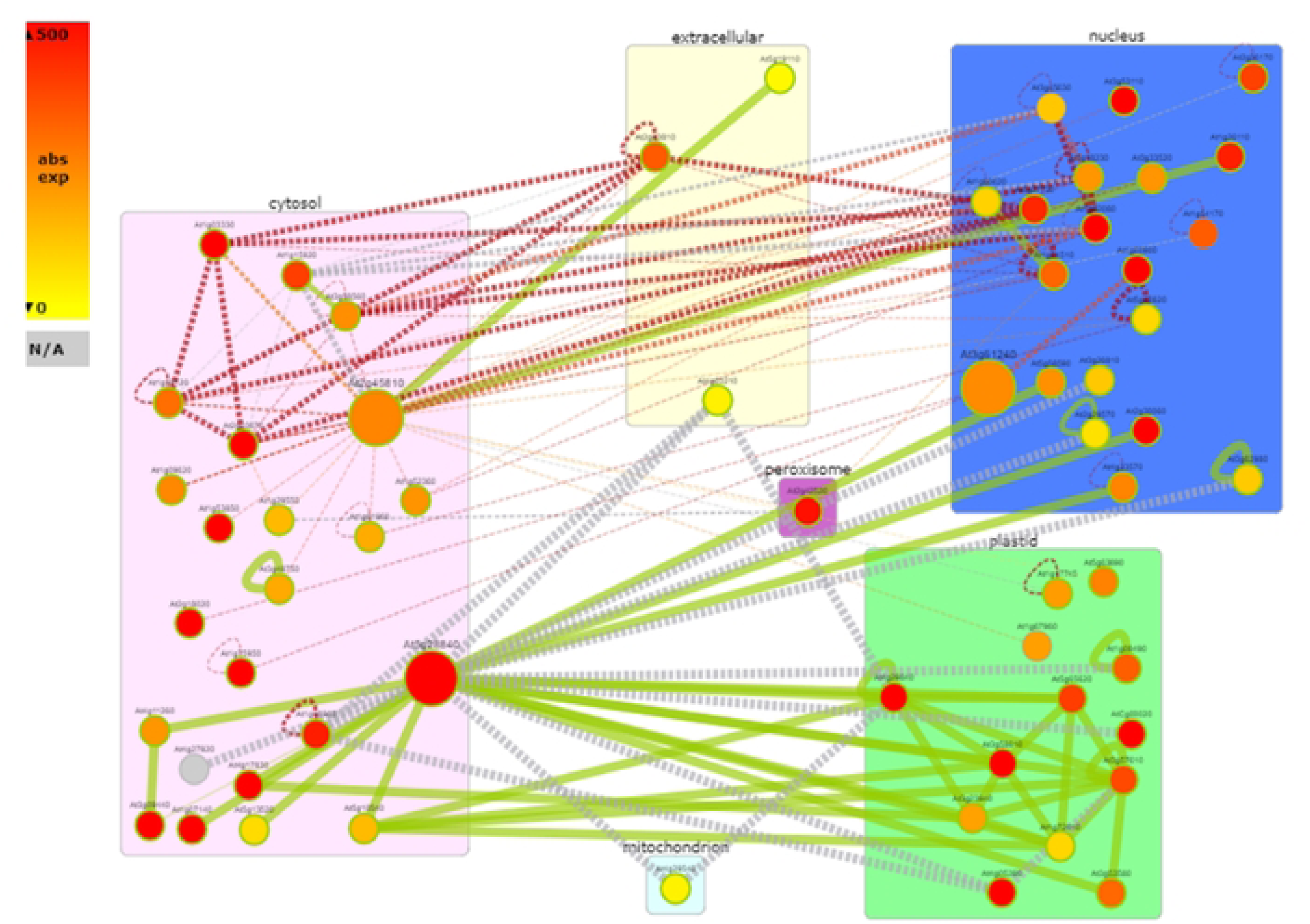
Protein-protein interactors of AT2G45810 and its interactors, AT5G28840 and AT3G61240, with expression levels overlaid in guard cells plus 100 um ABA.

## Conclusion

Our knowledge of plant development, physiology, and genetics has greatly improved thanks to Arabidopsis. Arabidopsis will continue to be important in future plant biotechnology research in several areas, including plant-microbe interactions, gene editing, metabolic engineering, understanding plant responses to environmental stress, crop improvement, biofuel production, etc. The transcriptomic analysis demonstrated the complex regulatory mechanisms governing important developmental transitions and stress responses, as well as the dynamic character of the Arabidopsis transcriptome and comparative genomics. The discovery of a large number of genes with variable expression has paved the way for our comprehension of the molecular underpinnings of many physiological processes, from flower formation to seed germination. A deeper comprehension of plant development and adaptability is made possible by the identification of important regulatory genes and signaling pathways that are involved in these processes. Transcriptome and developmental analyses in *Arabidopsis thaliana* cooperate synergistically to provide a more thorough understanding of plant biology. To learn more about the molecular mechanisms behind plant development and stress responses, future research projects should make use of cutting-edge technology and interdisciplinary techniques. The consequences of our findings will open up an excess of a specific gene expression pattern, along with information from embryonic developmental stages, maps, and DNA damage analysis.

## Acknowledgment of Gratitude

The authors would like to express the most profound gratitude to Department of Agriculture, Bangabandhu Sheikh Mujibur Rahman Science and Technology University, Gopalganj-8100.

## Author Contributions

Conceptualization, ME; Data curation - figure, diagram, and map, ME, AH and ZAJ; Data curation – table, ME, AH and ZAJ; Formal analysis, ME, AH, SKB, ZAJ and MSA; Investigation, ZAJ and MSA; Methodology, ME, AH, SKB, ZAJ and MSA; Project administration, ZAJ and MSA Resources, ME, AH and SKB; Software, ME and AH; Supervision: ZAJ and MSA Validation, ME, AH, SKB, ZAJ and MSA Visualization, ME, AH, SKB, ZAJ and MSA Writing - original manuscript, ME, AH, SKB, ZAJ and MSA; Writing - review and editing, ME, AH, SKB, ZAJ and MSA;

